# Evidence that humans underestimate body mass in microgravity: kinematic signatures in reaching movements during spaceflight

**DOI:** 10.1101/2025.05.12.653611

**Authors:** Zhaoran Zhang, Yu Tian, Chunhui Wang, Changhua Jiang, Bo Wang, Hongqiang Yu, Rui Zhao, Kunlin Wei

## Abstract

Astronauts consistently exhibit slower movements in microgravity, even during tasks requiring rapid responses. The sensorimotor mechanisms underlying this general slowing remain debated. Two hypotheses have been proposed: either the sensorimotor system adopts a conservative control strategy for safety and postural stability, or the system underestimates body mass due to reduced inputs from proprioceptive receptors. To dissociate these opinions, we studied twelve taikonauts aboard the China Space Station performing a classical hand-reaching task. Compared to their pre-flight performance and to an age-matched control group, participants showed increased movement durations and altered kinematic profiles in microgravity. Model-based analyses of motor control parameters revealed that these changes stemmed from reduced initial force generation in the feedforward control phase followed by compensatory feedback-based corrections. These findings providing support for the body mass underestimation hypothesis while being inconsistent with the strategic slowing hypothesis. Importantly, the sensory estimate of bodily property in microgravity is biased but immune from sensorimotor adaptation, calling for an extension of existing theories of motor learning.

## Introduction

Successful space exploration fundamentally depends on efficient sensorimotor control under microgravity conditions, where both movement accuracy and speed are critical (Strangman et al., 2014). A consistently observed phenomenon is that astronauts exhibit slower movements during routine activities in space stations (Kubis et al., 1977). This general movement slowing persists even in controlled experimental settings where astronauts are securely fastened without concerns for postural stability or safety. Indeed, fast movements in microgravity can be disadvantageous, potentially destabilizing posture, complicating the arrest of accelerated body segments, and increasing collision risks in confined spacecraft environments (Bock, 1998; Layne et al., 2001; Paloski et al., 1999; Reschke et al., 1998; Tafforin et al., 1989). Yet even when body stability is ensured, well-practiced goal-directed actions consistently show prolonged movement durations across various tasks, from hand pointing to visual targets (Berger et al., 1997; Bock et al., 2001, 2003; Mechtcheriakov et al., 2002) to step tracking with joysticks (Newman & Lathan, 1999; Sangals et al., 1999). This slowing manifests as reduced peak speeds in hand pointing (Mechtcheriakov et al., 2002; Sangals et al., 1999; Weber & Stelzer, 2022). Despite extensive documentation of this phenomenon, the underlying sensorimotor mechanisms remain unclear (Bock et al., 1996b; Weber et al., 2019), presenting a significant challenge for optimizing human performance in space.

Two primary hypotheses have emerged to explain the slowing of goal-directed actions in microgravity. The conservative control hypothesis suggests that the sensorimotor system adopts a generalized strategy prioritizing safety and postural stability over speed (Berger et al., 1997; Bock et al., 2010; Mechtcheriakov et al., 2002). This adaptation may arise from the novel challenges of movement control in microgravity, where even simple actions can propagate unexpected forces through body segments. The persistent application of this conservative strategy, as a generalized adaptation to the novel environment, could explain the movement slowing in microgravity. Alternatively, the body mass underestimation hypothesis builds on observations that humans consistently underestimate the mass of handheld objects in microgravity (Ross & Reschke, 1982), suggesting that similar perceptual biases might extend to the estimation of body segment properties (Bock et al., 1996b). Such misperception would directly impact the feedforward control of movement, potentially leading to systematic under-actuation of initial movements (Elliott et al., 2010, 2017).

Both hypotheses predict reduced peak speed and acceleration in microgravity, but they differ in their predictions about finer-grained movement kinematics. First, typical reaching movement exhibits a symmetrical bell-shaped speed profile, which minimizes energy expenditure while maximizing accuracy or smoothness according to optimal control principles (Flash & Hogan, 1985; Todorov, 2004). The conservative control hypothesis suggests a maintained temporal symmetry as longer durations are strategically planned for optimal performance, the peak speed and acceleration are thus delayed in time (Figure 1A). In contrast, the body mass underestimation hypothesis predicts that insufficient initial force generation, resulting from a systematic sensory bias in internal models that predict movement dynamics, would lead to earlier occurrence of peak speed and acceleration (Elliott et al., 2010, 2017). Previously, the timing of these peaks has shown mixed results, with some studies reporting earlier peaks (Sangals et al., 1999), others finding delays (Fowler et al., 2008), and some showing no temporal shifts (Berger et al., 1997; Mechtcheriakov et al., 2002). Second, the hypotheses make distinct predictions about feedback-based corrections during late reaching. The conservative control hypothesis suggests that movements would remain well-planned with near-optimal execution, thus requiring minimal corrections. Instead, the mass underestimation hypothesis predicts that initial underactuation would necessitate subsequent feedback-based corrections to reach the target (Chua & Elliott, 1993; Smeets & Brenner, 1995). These corrections would manifest as additional submovements, contributing to asymmetric speed profiles (Figure 1A). Current evidence for increased feedback-based corrections in microgravity remains inconsistent: while some studies support their presence (Sangals et al., 1999), others show conflicting results (Mechtcheriakov et al., 2002) or opposite effects (Fowler et al., 2008). If present, these corrective submovements should predict movement duration prolongation, serving as important evidence favoring the mass-underestimation account for the general slowing in microgravity.

**Figure 1.**
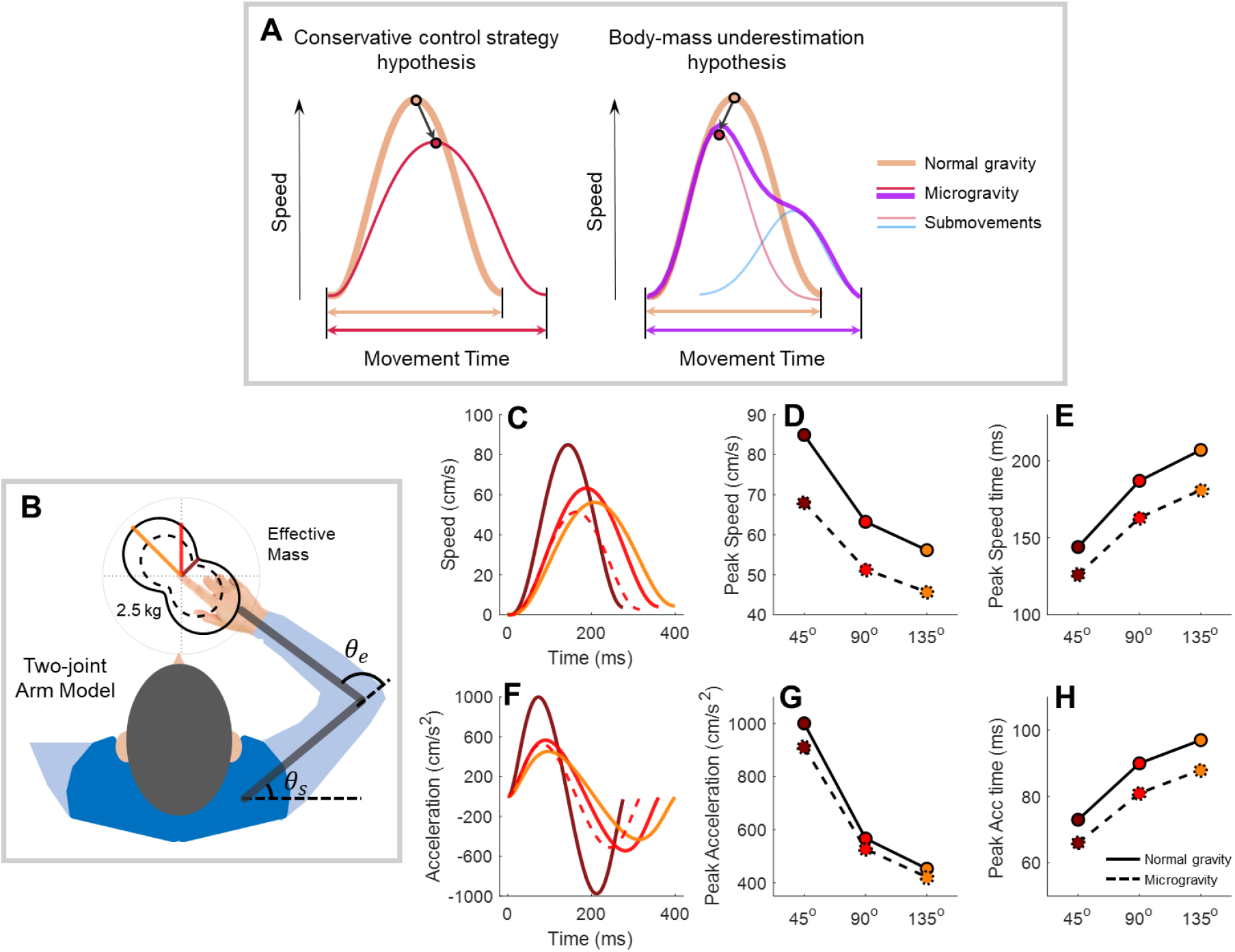
Model simulations revealing characteristic kinematic features of reaching movements. **A)** Speed profile predictions under two hypotheses. Under normal gravity, hand speed follows a typical profile (bold apricot lines). According to the body mass underestimation hypothesis (left panel), initial underactuation produces a lower, earlier peak speed (light red), necessitating a corrective submovement (light blue) that results in an asymmetric, prolonged profile (purple). The conservative strategy hypothesis (right panel) predicts symmetric slowing with delayed peak speed (red). **B)** Two-joint arm model and its simulated effective mass. The spatial distribution of effective mass is shown by black curves (solid: normal gravity; dashed: 30% mass underestimation in microgravity). Colored lines indicate effective mass values for our three experimental target directions (45°, 90°, 135°). **C & F)** Representative speed and acceleration profiles simulated using effective masses from B. Profiles shown for normal (solid) and underestimated (dashed) body mass in the 90° direction demonstrate reduced, earlier peaks with mass underestimation. **D-E & G-H)** Simulated kinematic parameters across movement directions under normal (solid) and underestimated (dashed) mass conditions. Mass underestimation consistently leads to reduced amplitude and earlier timing of speed/acceleration peaks across all directions. Model simulation details are provided in the Methods.

To help distinguish between these theories, we leveraged a unique biomechanical property of arm movements: the natural variation in effective limb mass across movement directions, known as anisotropic inertia (Mussa-Ivaldi et al., 1985). Based on a two-joint arm model (Li & Todorov, 2004), reaching movements toward targets in the frontal-medial direction involve greater effective mass compared to those in the frontal-lateral direction. Based on the motor utility theory, both the magnitude and timing of peak speed and acceleration should be systematically modulated by target direction (Shadmehr et al., 2016). Similarly, if astronauts underestimate their limb mass by approximately 30% in microgravity (Crevecoeur et al., 2014), we would expect reduced peak speed/acceleration and earlier occurrence of these peaks across all the directions. Here we examined twelve taikonauts before, during, and after their 3- or 6-month missions on the China Space Station (CSS) and compared their performance to age-matched ground controls. Participants performed rapid reaching movements to targets specifically chosen to engage different effective masses. Our approach allows us to dissociate the effects of mass underestimation from strategic control by examining how movement kinematics vary with direction-dependent effective mass.

## Results

We compared the performance of 12 taikonauts to that of age-matched ground controls across multiple test sessions (Figure 6B and Table S1). Participants performed rapid reaching movements toward one of three targets on a tablet, presented in pseudorandom order to minimize movement automation and ensure active planning for each reach. Participants were instructed to reach the target with both speed and accuracy, pausing briefly before initiating the return. The emphasized requirement of moving fast would critically test whether movement slowing is ubiquitous in microgravity. The three target directions (45°, 90°, and 135°) were chosen to vary the effective mass of the moving limb, as verified through biomechanical simulations based on the two-joint arm model ((Li & Todorov, 2004); Figure 1B). Our model simulations not only demonstrated the direction-dependent variation in limb mass but also predicted how actual and misperceived limb mass would affect reaching kinematics (Figure 1C-H). These predictions were derived by combining the movement utility framework for estimating planned movement duration ((Shadmehr et al., 2016); Figure S1A-B) with a forward optimal controller for simulating reaching trajectories ((Todorov, 2005); Figure 1-figure supplement 1C-D). Notably, the model predicted that underestimating limb mass would lead to reduced and earlier peak speed/acceleration across directions (Figure 1C-H).

### Movement Accuracy and Reaction Time Remain Intact in Microgravity

All participants successfully completed the task. Invalid trials, comprising early initiation (RT < 150 ms), late initiation (RT > 400 ms), or measurement failures, accounted for only 1.87% of trials in the experimental group and 0.70% in the control group. The percentage of invalid trials remained stable across the pre-in and post-flight phases (Friedman’s tests, all *p* > 0.1).

Reaching accuracy remained high despite the stringent speed requirement of completing movements within 650 ms after target appearance (Figure 2-figure supplement 2). Analysis of endpoint error using a 3 (target direction) × 3 (phase) repeated-measures ANOVA revealed no significant differences across phases for either group. While the experimental group showed slightly better accuracy for the 45° target, this difference was minimal (0.04 cm over a 12 cm reach distance).

**Figure 2.**
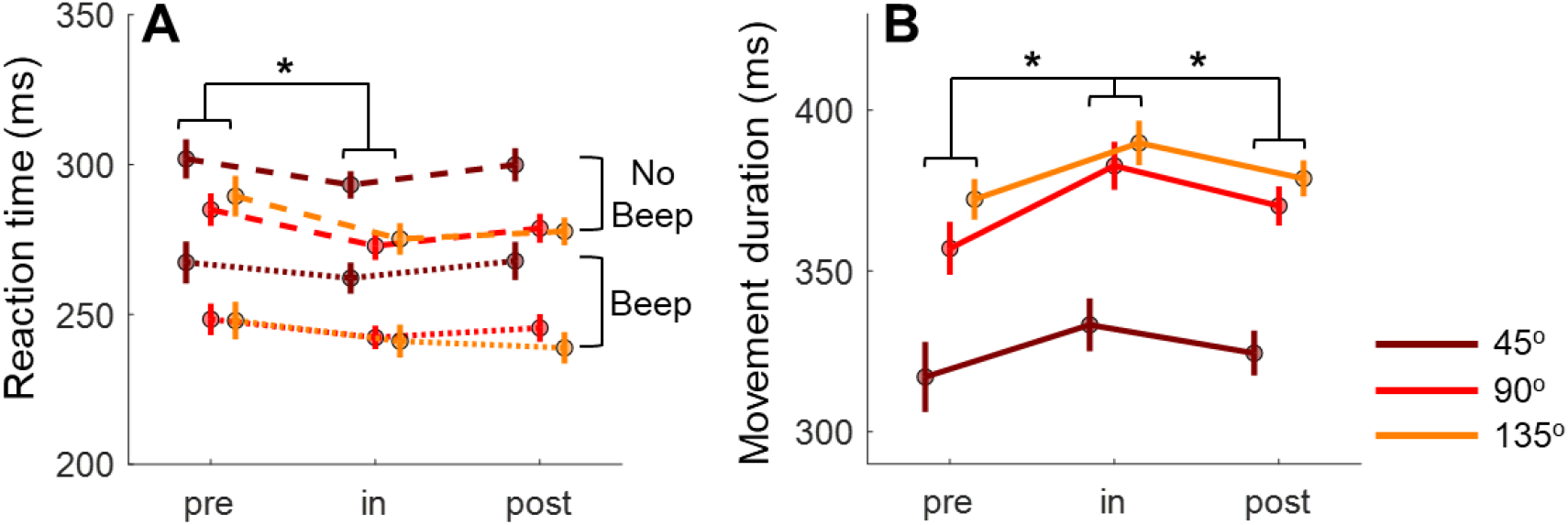
Reaction time and movement duration in the experimental group. **A)** Median reaction times plotted by experimental phase (x-axis) and target direction (different colors). Dashed lines and dot lines represent the “beep” and “no beep” conditions. **B)** Average movement durations shown in the same format. Data are combined across beep conditions due to no significant differences between them. In both panels, error bars denote standard error across participants. Asterisks indicate significance levels for post-hoc phase comparisons (\**p* < 0.05, \*\**p* < 0.01, \*\*\**p* < 0.001).

To introduce sufficient variability in reaction times for assessing speed–accuracy trade-offs (Sutter et al., 2021), a beep sound was played upon the target appearance in half of the trials. The random auditory signal tended to accelerate responses, thereby widening the distribution of reaction times in our dataset. The beep effectively reduced median reaction times by approximately 38 ms (242 ± 49 ms vs. 281 ± 50 ms; Figure 2A; see control group data in Figure 2-figure supplement 1A). To evaluate microgravity effects, we pooled data from both beep conditions and conducted a 3 (direction) × 3 (phase) repeated-measures ANOVA. The experimental group showed significant main effects of direction (F(2,22) = 25.260, *p* < 0.001, partial η^2^ = 0.697) and phase (F(2,22) = 5.820, *p* = 0.014, partial η^2^ = 0.346), without interaction (F(4,44) = 2.246, *p* = 0.111). Post-hoc analyses revealed slower reaction times for the 45° direction compared to both 90° (*p* < 0.001, d = 1.473) and 135° (*p* = 0.003, d = 1.427). Contrary to the conservative-strategy hypothesis, taikonauts did not show generalized slowing in reaction time; they actually had faster reaction times during spaceflight, incompatible with a generalized slowing strategy. Reactions were faster during the in-flight phase compared to pre-flight (*p* = 0.037, d = 0.803), with no significant difference between in-flight and post-flight phases (*p* = 0.127). Similar effects were observed in the control group (Figure 2-figure supplement 1A), suggesting that the reduced reaction times likely reflect practice effects rather than microgravity influence. Thus, participants maintained, and even improved, their ability to rapidly initiate movements in microgravity with reaching accuracy.

### Microgravity leads to slower movement

Although showing no decrement in movement accuracy or RT, the taikonauts exhibited prolonged movement durations (MD) during the in-flight phase compared to both pre- and post-flight phases (Figure 2B). A two-way repeated-measures ANOVA reveals significant main effects of direction (F(2,22) = 62.555, *p* < 0.001, partial η^2^ = 0.850) and phase (F(2,22) = 6.761, *p* = 0.015, partial η^2^ = 0.381), with no significant interaction (F(4,44) = 1.413, *p* = 0.263). Post-hoc comparisons showed that MD was significantly prolonged during the in-flight phase (vs. pre-flight: *p* = 0.014, d = 0.866; vs. post-flight: *p* = 0.013, d = 0.472). MD was also significantly greater for the 90° and 135° directions—those associated with larger effective mass—compared to the 45° direction (all p < 0.001 in pairwise comparisons). In contrast, the control group showed a similar directional effect but, importantly, did not show a phase effect (Figure 2—figure supplement 1), suggesting that movement slowing is specific to spaceflight.

### Microgravity leads to reduced peak acceleration/speed and advanced timing of these peaks

The initial movement exhibited signs of underactuation during spaceflight, consistent with the body mass underestimation hypothesis (Figure 2—figure supplement 6). Peak acceleration and peak speed, established measures of feedforward control (Elliott et al., 2010, 2017), both showed significant decreases during the in-flight phase (Figure 3). For peak acceleration, there were significant main effects of phase (F(2,22) = 6.401, *p* = 0.015, partial η^2^ = 0.368) and direction (F(2,22) = 75.516, *p* < 0.001, partial η^2^ = 0.873), along with an interaction effect (F(4,44) = 4.548, *p* = 0.016, partial η^2^ = 0.293). Consistent with model simulations, the 45° direction, associated with a lower effective mass, showed significantly higher peak acceleration than the other two directions (Figure 3A; all p < 0.001 in pairwise comparisons). Peak acceleration decreased significantly during spaceflight across all three directions (simple main effects: 45°, *p* = 0.020, η^2^ = 0.543; 90°, *p* = 0.004, η^2^ = 0.673; 135°, *p* = 0.005, η^2^ = 0.657). Planned contrasts further indicated lower peak acceleration in the in-flight phase compared to pre-flight (45°, *p* = 0.006, η^2^ = 0.516; 90°, *p* = 0.008, η^2^ = 0.485; 135°, *p* = 0.003, η^2^ = 0.560), with marginal recovery in the post-flight phase for two directions (45°, *p* = 0.071, η^2^ = 0.267; 90°, *p* = 0.062, η^2^ = 0.282) and significant recovery for one direction (135°, *p* = 0.029, η^2^ = 0.363). In contrast, the control group displayed only the direction effect (Figure 3—figure supplement 1A-B), confirming that the underactuation was specific to microgravity.

**Figure 3.**
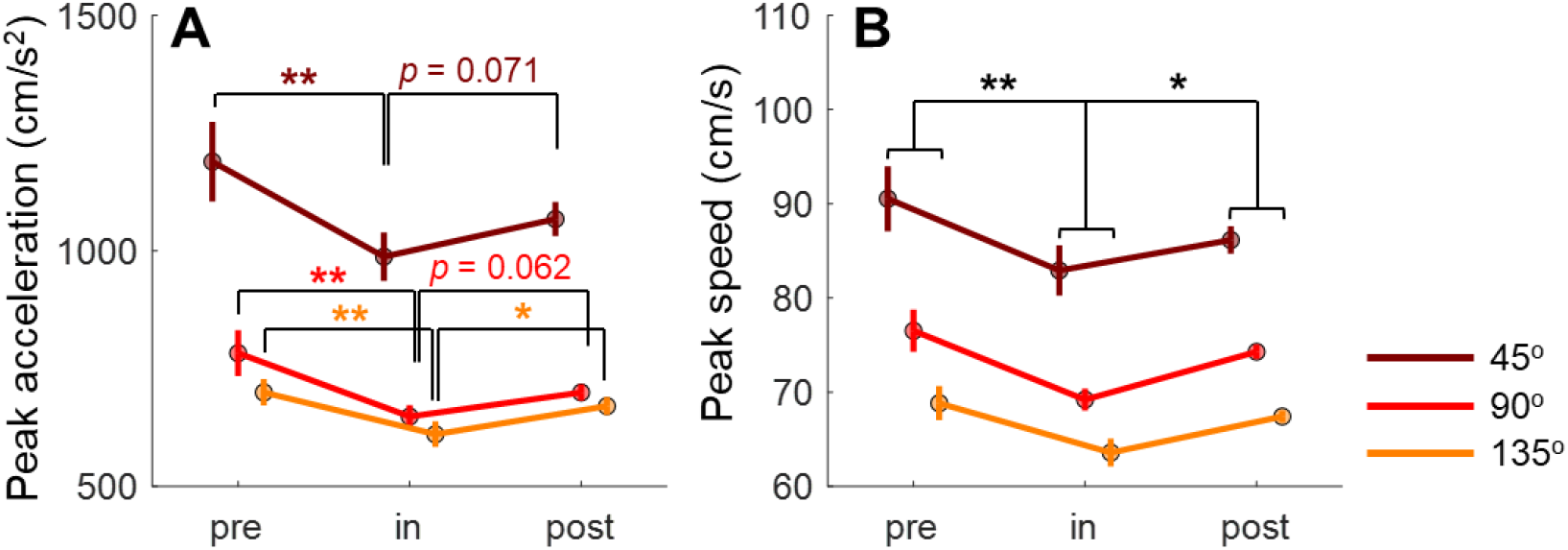
Magnitude changes of peak acceleration and speed during spaceflight. **A)** Peak acceleration across experimental phases and movement directions, showing systematic decreases during the in-flight phase. **B)** Peak speed data presented in the same format, demonstrating parallel changes to peak acceleration. Error bars indicate standard error across participants. Statistical significance levels are shown for both overall phase comparisons (black asterisks) and planned contrasts within each direction (colored asterisks): \**p* < 0.05, \*\**p* < 0.01, \*\*\**p* < 0.001.

Peak speed showed parallel changes during spaceflight (Figure 3B). Analysis revealed significant main effects of phase (F(2,22) = 7.043, *p* = 0.009, partial η^2^ = 0.390) and direction (F(2,22) = 114.420, *p* < 0.001, partial η^2^ = 0.912), without interaction (F(4,44) = 1.929, *p* = 0.161). Peak speed was significantly reduced during the in-flight phase compared to both pre-flight (*p* = 0.004, d = 0.879) and post-flight phases (*p* = 0.046, d = 0.529), with evidence of recovery post-flight (pre-vs. post-flight: *p* = 0.483, d = 0.349). Target direction systematically modulated peak speed (all pairwise comparisons, *p* < 0.001), with the 45° direction showing highest speeds and 135° lowest—matching predictions based on the movement utility theory. The control group showed only direction effects without phase-related changes (Figure 3—figure supplement 1B), confirming the specificity of these changes to microgravity exposure.

The critical test between the two hypotheses of movement slowing lies in whether peak acceleration and peak speed occur earlier or later in microgravity. The body mass underestimation hypothesis predicts earlier timing, whereas the conservative control strategy hypothesis predicts delayed timing. We observed an overall timing advance of these two measures. For peak acceleration time in the experimental group (Figure 4A), two-way repeated-measures ANOVAs revealed significant main effects of phase (F(2,22) = 5.189, *p* = 0.024, partial η^2^ = 0.321) and direction (F(2,22) = 30.375, *p* < 0.001, partial η^2^ = 0.734), along with an interaction effect (F(4,44) = 4.503, *p* = 0.013, partial η^2^ = 0.290). In line with predictions, peak acceleration appeared significantly earlier in the 45° direction than other directions (45° vs. 90°, *p* < 0.001, d = 1.675; 45° vs. 135°, *p* < 0.001, d = 1.491). The phase effect emerged only for directions with higher effective mass (simple main effects, 90°: F(2,10) = 6.660, *p* = 0.015, η^2^ = 0.571; 135°: F(2,10) = 8.411, *p* = 0.007, η^2^ = 0.627). Planned contrasts confirmed earlier peak acceleration during spaceflight compared to post-flight (90°, *p* = 0.005, η^2^ = 0.530; 135°, *p* = 0.004, η^2^ = 0.537), with marginal differences versus pre-flight (135°, *p* = 0.065). The control group showed only directional effects without phase effect (Figure 4—figure supplement 1A).

**Figure 4.**
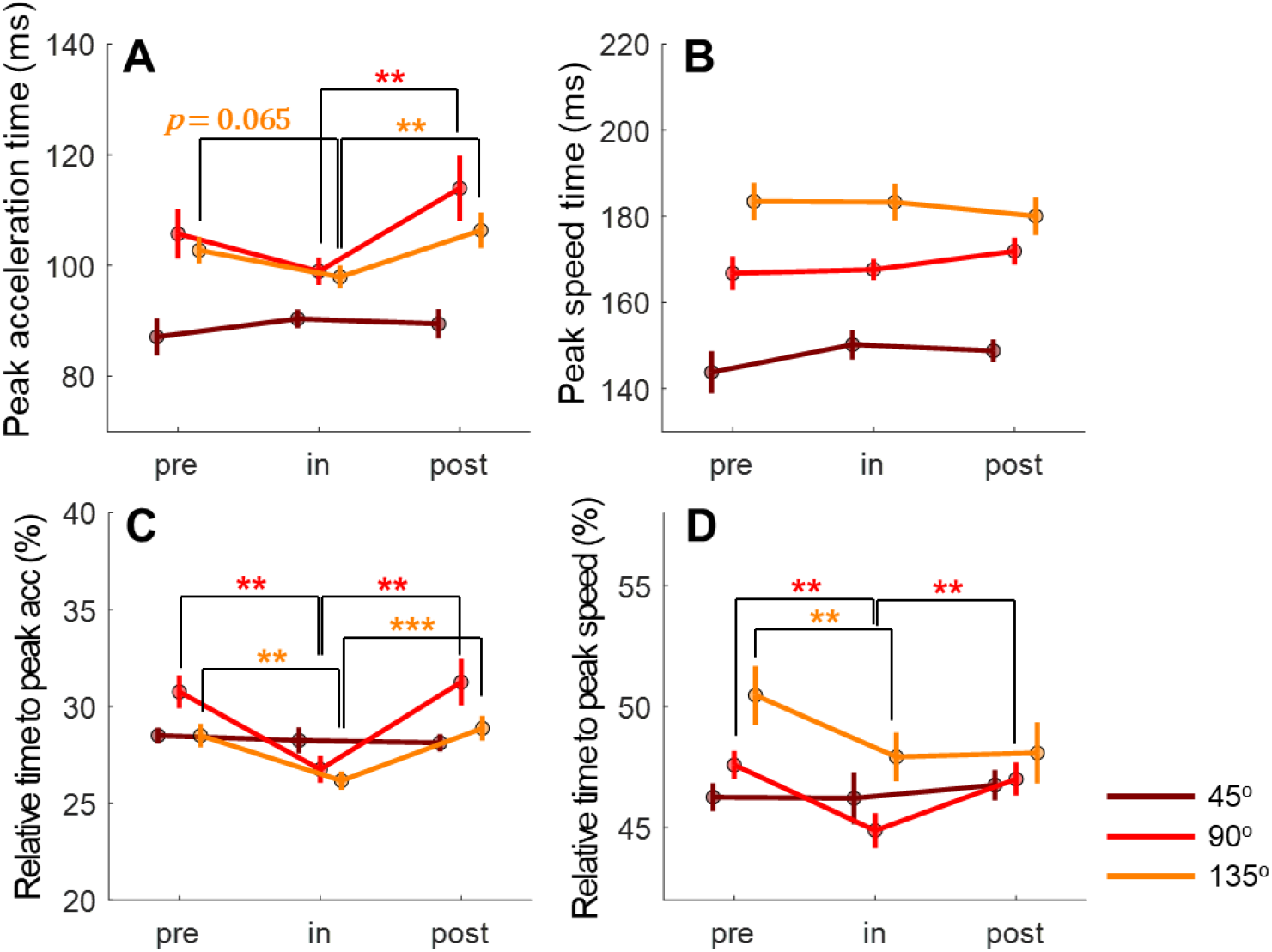
Temporal analysis of peak acceleration and peak speed during spaceflight. **A-B)** Time to peak acceleration and peak speed across experimental phases and movement directions. **C-D)** Relative timing analysis showing these same peaks as a percentage of total movement duration, revealing more pronounced microgravity effects than absolute timing measures. Error bars indicate standard error across participants. Asterisks denote significance levels for planned phase comparisons (\**p* < 0.05, \*\**p* < 0.01, \*\*\**p* < 0.001). Note the systematic modulation of timing by phase, particularly for high effective mass conditions.

Given that overall movement duration is prolonged in microgravity, a more direct way to assess the symmetry of the speed profile is to compute the relative timing of speed/acceleration peaks as a percentage of MD. Indeed, the relative timing of peak acceleration showed the same reduction in microgravity with a more pronounced effect than absolute timing (Figure 4C). A two-way repeated-measures ANOVA revealed significant main effects of phase (F(2,22) = 15.414, *p* < 0.001, partial η^2^ = 0.584) and interaction (F(2,22) = 7.223, *p* = 0.002, partial η^2^ = 0.396), while the main effect of direction was marginally significant (F(2,22) = 3.374, *p* = 0.053, partial η^2^ = 0.235). Phase effects were significant for directions with higher effective mass (simple main effects: 90°, F(2,10) = 13.327, *p* = 0.002, η^2^ = 0.727; 135°, F(2,10) = 24.574, *p* < 0.001, η^2^ = 0.831) but not for 45° (F(2,10) = 0.609, *p* = 0.563). Planned contrasts confirmed significantly earlier timing during spaceflight compared to both pre- and post-flight phases (all p < 0.002 for 90° and 135° directions). None of these effects appeared in the control group (Figure 4—figure supplement 1C).

Peak speed also showed a timing advance, but only for relative timing. For absolute timing (Figure 4B), a two-way repeated-measures ANOVA revealed a significant main effect of direction (F(2,22) = 57.428, *p* < 0.001, partial η^2^ = 0.839) and a marginal interaction (F(4,44) = 2.233, *p* = 0.095, partial η^2^ = 0.169), without a phase effect (F(2,22) = 0.093, *p* = 0.864). In contrast, relative timing analysis (Figure 4D) showed significant effects of phase (F(2,22) = 8.941, *p* = 0.004, partial η^2^ = 0.448), direction (F(2,22) = 4.991, *p* = 0.030, partial η^2^ = 0.312), and their interaction (F(4,44) = 4.755, *p* = 0.015, partial η^2^ = 0.302). The phase effect appeared only for directions with higher effective mass (90°: F(2,10) = 17.278, *p* = 0.001, η^2^ = 0.776; 135°: F(2,10) = 6.189, *p* = 0.018, η^2^ = 0.553). Planned contrasts revealed significantly earlier relative timing during spaceflight versus pre-flight for both 90° (*p* < 0.001, η^2^ = 0.770) and 135° (*p* = 0.007, η^2^ = 0.496) directions, with significant post-flight recovery in 90° (*p* = 0.003, η^2^ = 0.575). The control group showed only directional effects (Figure 4—figure supplement 1B & 1D). Thus, after accounting for the prolongation of movement duration, we observed a temporal advance of peak acceleration and speed during spaceflight.

### Movement slowing is associated with corrective submovements

The feedforward component of movement, captured by our model simulations, accounts for only the initial phase of reaching. With peak acceleration and speed occurring at approximately 80 ms and 150 ms after movement onset, and total movement duration spanning 300-400 ms, participants had ample opportunity for feedback-based corrections (Dimitriou et al., 2013). These corrections, if present, would manifest as secondary submovements (D. E. Meyer et al., 1988). While optimal reaching movements typically show symmetrical speed profiles (Flash & Hogan, 1985), the presence of corrective submovements creates characteristic right-skewed speed profiles (Elliott et al., 2010; D. E. Meyer et al., 1988; Woodworth, 1899). To identify such corrections, we employed an established decomposition method (Rohrer & Hogan, 2006) to detect trials containing both primary and secondary submovements (two-peak trials; Figure 5A).

**Figure 5.**
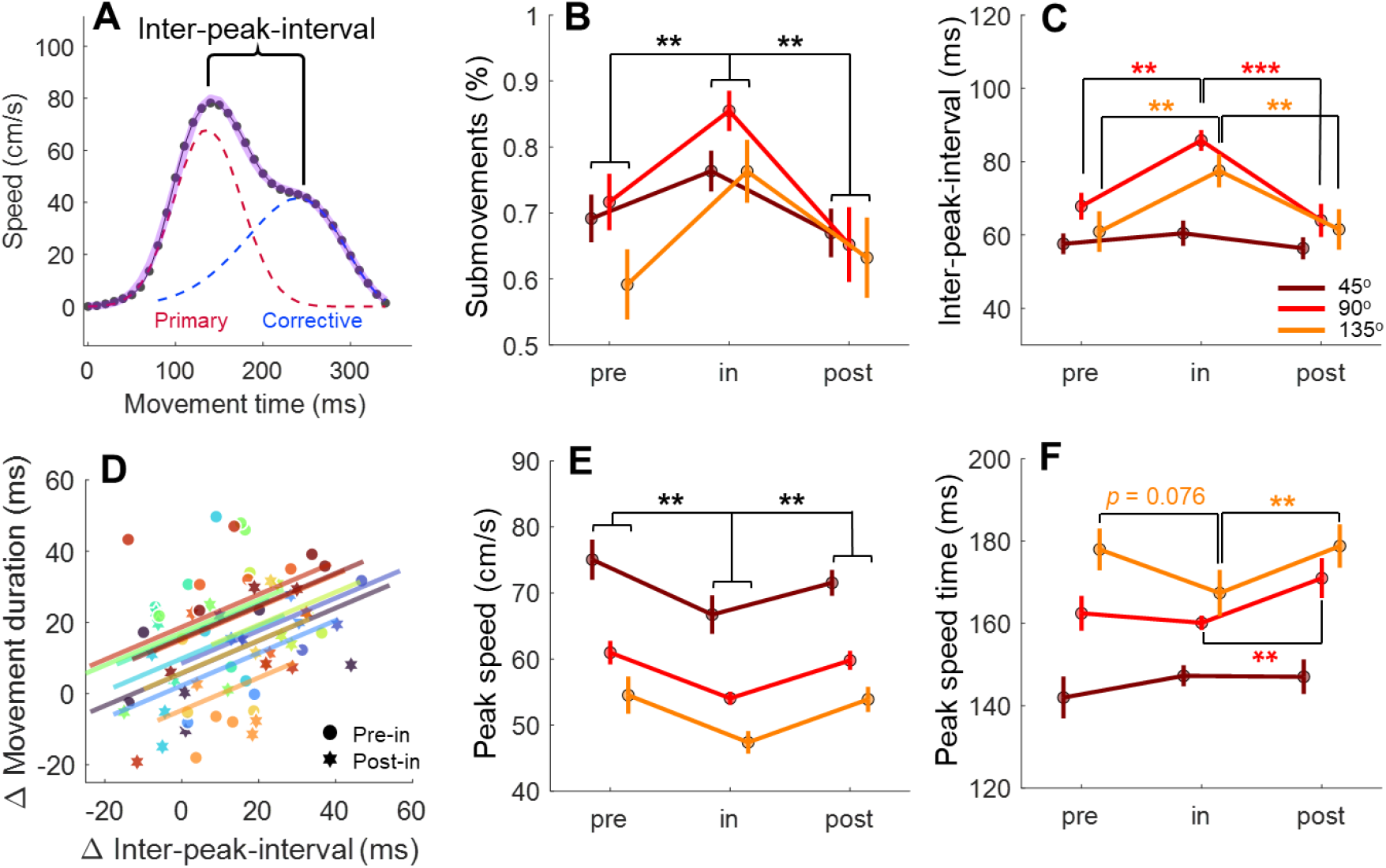
Submovements analysis revealed changes in corrective movements during spaceflight. **A)** Decomposition of a representative two-peak speed profile, illustrating primary and corrective submovements separated by the inter-peak interval (IPI). **B)** Proportion of movements showing corrective submovements across phases and directions. **C)** Magnitude of feedback corrections quantified by IPI, showing direction-dependent increases in microgravity. **D)** Linear relationship between feedback correction changes (ΔIPI) and movement slowing (ΔMD). Each participant (shown in different colors) contributed multiple data points from different phases and directions. **E-F)** Primary submovement characteristics: peak speed amplitude and timing. Error bars denote standard error across participants; asterisks indicate significance (\**p* < 0.05, \*\**p* < 0.01, \*\*\**p* < 0.001).

Analysis revealed a striking increase in corrective movements during spaceflight. The percentage of trials showing two distinct submovements increased significantly during the in-flight phase (Figure 5B), evidenced by a main effect of phase (F(2,22) = 13.969, *p* < 0.001, partial η^2^ = 0.559) and a marginal phase-by-direction interaction (F(2,22) = 2.642, *p* = 0.054, partial η^2^ = 0.194). This effect was specific to microgravity exposure, as the control group showed no significant changes across phases (all *p* > 0.05; Figure 5—figure supplement 1A). Notably, the increase in corrective movements was most pronounced for directions with higher effective mass (simple main effects: 90°, F(2,10) = 11.426, *p* = 0.003, η^2^ = 0.696; 135°, F(2,10) = 7.292, *p* = 0.011, η^2^ = 0.593).

To understand how feedback corrections contribute to movement slowing, we analyzed the inter-peak interval (IPI) between primary and corrective submovements, an established measure of feedback-based control (Craik, 1947; D. Meyer et al., 2018; D. E. Meyer et al., 1988). The taikonauts showed markedly longer IPIs during spaceflight (Figure 5C), with a two-way ANOVA revealing significant effects of phase (F(2,22) = 11.150, *p* = 0.001, partial η^2^ = 0.503), direction (F(2,22) = 4.721, *p* = 0.026, partial η^2^ = 0.300), and their interaction (F(2,22) = 6.559, *p* = 0.002, partial η^2^ = 0.374). Critically, IPI increases were confined to directions with greater effective mass (90°, F(2,10) = 11.573, *p* = 0.002, η^2^ = 0.698; 135°, F(2,10) = 13.371, *p* = 0.001, η^2^ = 0.728). The control group exhibited only directional effects (F(2,22) = 19.277, *p* < 0.001, partial η^2^ = 0.637; Figure 5—figure supplement 1B) without phase-related changes (all *p* > 0.13).

To establish whether these enlarged corrective movements explain the overall movement slowing, we examined how changes in IPI (ΔIPI) predict changes in movement duration (ΔMD) during spaceflight (Figure 5D). A Linear Mixed Model analysis incorporating ΔIPI, phase transition (pre-to-in and post-to-in), and direction revealed that ΔIPI significantly predicted ΔMD (β = 0.458 [0.144, 0.771], t(67) = 2.915, *p* = 0.005, partial-R^2^ = 0.103). Phase transition also showed significance (β = −0.01 [−0.017, −0.001], t(67) = −2.696, *p* = 0.009, partial-R^2^ = 0.086), with stronger effects for pre-to-in versus post-to-in transitions, indicating incomplete recovery post-flight. Movement direction showed no significant effect (all *p* > 0.42), suggesting that the relationship between corrective movements and duration slowing generalizes across reaching directions.

The analysis of primary submovements revealed even stronger evidence for feedforward control changes in microgravity. By focusing on just the primary submovement in two-peak trials, we found more pronounced effects than when analyzing the overall movement (comparing Figure 5E to Figure 5—figure supplement 1C). The primary submovement showed a robust reduction in peak speed during spaceflight (main effect of phase: F(2,22) = 10.363, *p* = 0.001, partial η^2^ = 0.485) that was consistent across directions (interaction: F(4,44) = 0.382, *p* = 0.711; Figure 5E). More tellingly, while the overall movement showed timing changes only in relative measures, the primary submovement exhibited earlier peak speed timing in absolute terms (phase-by-direction interaction: F(2,22) = 4.264, *p* = 0.012, partial η^2^ = 0.279). This temporal advance was most evident in directions with higher effective mass (planned contrasts vs. post-flight: 135°: *p* = 0.008, η^2^ = 0.615; 90°: *p* = 0.076, η^2^ = 0.403) but absent in the 45° direction (*p* = 0.188, η^2^ = 0.284). The selective nature of the primary submovement in both magnitude and timing (comparing Figure 4B and Figure 5F) provides compelling evidence for mass-dependent changes in feedforward control. The control group showed only the expected directional variations (F(2,22) = 46.259, *p* < 0.001, partial η^2^ = 0.808; Figure 5—figure supplement 1D) without any phase effects (all *p* > 0.13), confirming these changes as specific to microgravity exposure.

### The microgravity effects on movement kinematics are directionally dependent

Reaching movements are inherently directionally dependent due to anisotropies in limb dynamics (Gordon et al., 1994; Shadmehr et al., 2016). Even before microgravity exposure, key kinematic variables—peak speed, peak acceleration, and their timing—exhibited consistent directional differences, as shown in Figure 1C–H. This baseline direction-dependency suggests that effective mass varies across movement directions and influences movement execution under normal gravity. Building on this observation, we tested whether microgravity-induced changes would also follow a direction-specific pattern, as predicted by the mass underestimation hypothesis.

To assess microgravity-induced changes, we quantified the directional differences (Δ) in peak speed, peak acceleration, and their timing, comparing in-flight measurements to pre-flight baselines. The comparison between in-flight and post-flight was not included in this analysis since post-flight recovery was not complete (see peak speed and peak acceleration changes in Figure 3). As shown in Figure 6, these directional effects were evident across all kinematic metrics. For both Δ peak speed and Δ peak acceleration, the magnitude of change followed a consistent ranking across directions, with the largest effects at 45°, intermediate at 90°, and smallest at 135° (i.e., 45° > 90° > 135°). Similarly, for timing metrics—Δ peak speed time and Δ peak acceleration time—the shortest delays occurred at 45°, while 90° and 135° showed comparably larger shifts. Note the 45° did not show significant timing advance, as previously shown in Figure 5F.

**Figure 6.**
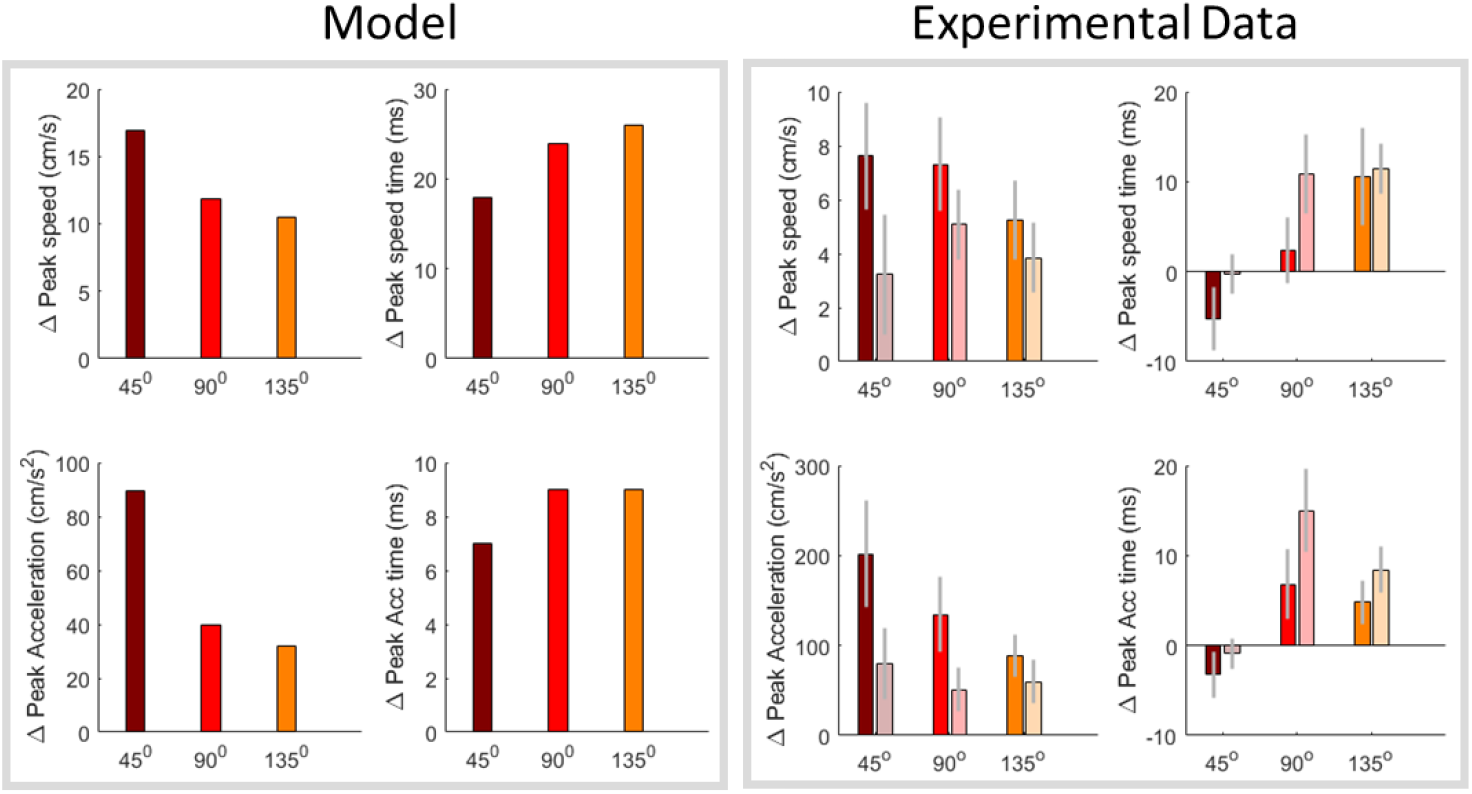
Direction-dependent effects of microgravity on peak kinematics and their timing. Left: model outputs. Right: experimental data shown as Δ relative to the in-flight session; solid bars = Δ(in − pre) and semi-transparent bars = Δ(in − post). Colors encode direction consistently across panels (e.g., 45° = darker hue, 90° = medium, 135° = lighter/orange). Panels (clockwise from top-left): Δ peak speed (cm/s), Δ peak speed time (ms), Δ peak acceleration time (ms), and Δ peak acceleration (cm/s^2^). Bars are group means; error bars denote standard error across participants.

Overall, the direction-dependent changes were roughly consistent with model predictions based on the mass underestimation hypothesis (Figure 6, comparing left and right panels). The model simulation similarly indicates that amplitude changes (Δ peak speed, Δ peak acceleration) and timing advances are rank ordered by movement directions. While the absolute magnitudes differed between model and data, the directional trends were broadly aligned. To assess the overall consistency between model predictions and experimental measurements, we performed repeated-measures correlation analyses between simulated and observed Δ values across directions (Bakdash & Marusich, 2017). Significant within-subject correlations were found for Δ peak speed time (*r*_*m*_ = 0.627, *p* < 0.001), Δ peak acceleration time (*r*_*m*_ = 0.591, *p* = 0.002), and Δ peak acceleration (*r*_*m*_ = 0.573, *p* = 0.003), while Δ peak speed did not reach significance (*r*_*m*_ = 0.334, *p* = 0.103). Together, these results suggest that the microgravity effect is direction-specific and the rank ordering of directions is broadly aligned with the predictions of the mass underestimation model. We note that these correlations evaluate the directional trend rather than the absolute magnitude of the effects; a precise quantitative match is not expected given the simplifications of the two-joint arm model.

### Speed-accuracy trade-offs remain intact during spaceflight

The speed-accuracy trade-off, a fundamental indicator of motor control efficiency, remained largely intact during spaceflight. We evaluated this trade-off using two complementary approaches. First, following the traditional Fitts’ framework =, we examined the relationship between movement duration and endpoint dispersion, which was not negatively affected by microgravity (Figure 2—figure supplement 3). Second, we analyzed the trade-off between reaction time and initial movement variance (Figure 2—figure supplement 4), which reflects the quality of movement planning (Sutter et al., 2021). This relationship persisted across all phases in both groups. Thus, while microgravity altered movement execution through mass underestimation, it did not fundamentally disrupt the sensorimotor system’s capacity to regulate the speed-accuracy trade-off. Additionally, in both groups, we found a significant negative correlation between movement duration (MD) and reaction time (RT), both across and within individuals (Figure 2—figure supplement 5). This finding indicates that participants moved faster when their RT was slower, and vice versa. This flexible motor adjustment, likely in response to the task requirement for rapid movements, remained consistent during spaceflight.

## Discussion

Long-term exposure to microgravity presents a unique environment that cannot be replicated on Earth, offering invaluable insights into human sensorimotor adaptation. Understanding how microgravity affects motor performance is crucial both for ensuring successful space exploration and for advancing our fundamental knowledge of motor control principles. Our study investigates the mechanisms underlying one of the most distinctive motor signatures in microgravity: the slowing of goal-directed actions during spaceflight. While taikonauts maintained reaching accuracy and rapid reaction times, their movements exhibited systematic changes in microgravity characterized by reduced peak speed and acceleration, together with direction-dependent advances in peak timing, particularly for movements involving higher effective mass. These persistent kinematic alterations (Figure 4—figure supplement 2) are not readily explained by a simple conservative-control account alone. Instead, they are consistent with an underactuation pattern that could arise, at least in part, from an underestimation of limb effective mass in microgravity.

These changes were broadly consistent with model predictions across movement directions with varying effective masses due to limb biomechanics, suggesting that the representation of body mass may affect movement control during spaceflight. Beyond the initial underactuation associated with altered feedforward control, we observed increased feedback-based corrective submovements that predicted movement duration prolongation in spaceflight. These effects were specific to microgravity exposure, occurring only during spaceflight with marked recovery post-flight, while ground controls showed no comparable changes. Notably, the taikonauts’ fundamental motor control capabilities remained largely intact, as evidenced by preserved movement accuracy, reaction time, and speed-accuracy trade-offs.

The underactuation observed in initial movements under microgravity extends findings from parabolic flight studies (Crevecoeur et al., 2014; Papaxanthis et al., 2005). When individuals performed reaching movements with handheld weights during parabolic flight-induced gravity alterations, their reaching trajectories and grip forces showed systematic changes suggesting mass misperception (Bock, 1998; Bock et al., 1996a, 1996b; Crevecoeur et al., 2014; Papaxanthis et al., 2005). In zero-gravity conditions, goal-directed arm movements were slower (Papaxanthis et al., 2005), and grip-load force coordination was altered (Crevecoeur et al., 2010, 2014), consistent with mass underestimation. Conversely, hypergravity (1.8g) induced increased peak acceleration and speed in reaching movements, along with elevated grip force during object manipulation, suggesting mass overestimation (Bock, 1998; Bock et al., 1992, 1996a, 1996b). Notably, modified grip-load force coordination reflects changes in feedforward control (Flanagan & Wing, 1997; Johansson & Edin, 1993), indicating that mass misestimation influences action planning. However, while parabolic flight studies capture only transient adaptations to rapid gravity transitions, our study targets general slowing during spaceflight and reveals the enduring effects of sustained microgravity exposure.

We also considered alternative explanations beyond mass underestimation. First, we tested whether a change in the cost-function of our optimal control model, which simulates a more conservative control, could account for movement slowing. This was implemented as a uniform rescaling of the state and control weights (Q and R) in the cost function (Crevecoeur et al., 2010). Increasing the scaling factors (α) will simulate the increase in control effort penalty and a more conservative control. However, while this manipulation reduced peak velocity and acceleration, it also delayed their occurrence, a pattern opposite to the observed effect. Thus, a uniform change in the cost function alone does not reproduce the observed combination of reduced peak amplitudes and advanced peak timing. Details of the cost-function simulations are provided in the Supplementary Materials (Supplementary Notes 1 and Figure 1—figure supplements 2 and 3). Second, changes in neuromuscular properties may also contribute to the observed underactuation. In microgravity, tonic muscle activity is likely reduced due to the absence of sustained antigravity demands, and descending vestibular inputs to motor neurons are diminished (Fisk et al., 1993). These changes could lower motor neuron excitability and alter muscle activation dynamics, potentially resulting in a weaker and slower initial response to an otherwise equivalent motor command—even in the absence of mass misestimation. While our simplified model treats the actuators as ideal torque generators and therefore does not capture these neuromuscular factors, such factors alone do not straightforwardly explain the direction-dependent pattern observed here. In the absence of additional direction-specific mechanisms, changes in tonic muscle activity, vestibular drive, or reflex gain would be expected to influence movement more generally across directions. The stronger effects observed in directions involving higher effective mass are therefore more naturally captured by a misrepresentation of inertial properties, although neuromuscular changes may also have contributed to the overall underactuation. Third, muscle weakness, a common effect during prolonged spaceflight, might also contribute to movement slowing. However, recent large-sample studies suggest that muscle strength reduction is mainly in lower-limb muscles that counter gravitational pull, not in upper-body muscles (Scott et al., 2023). Furthermore, the torques required in our task (~2 N·m for ~12 cm reaches) are relatively small, making it unlikely that muscle weakness is a primary factor of the observed kinematics, though a minor contribution cannot be excluded. Fourth, finger-screen friction might slow down the reaching movements, particularly in microgravity where astronauts must actively press on the screen to maintain contact in the absence of gravitational loading on the hand. However, during typical interaction with a touch screen, frictional forces are modest, generally ranging from 0.1 to 0.5 N (Ayyildiz et al., 2018), and directional variations are likely even smaller. These values are minimal compared to the 10–15 N required to accelerate the arm during reaching, making frictional anisotropy an unlikely source of the directional kinematic changes. Fifth, body stabilization in microgravity might also contribute to movement slowing in space. In fact, our participants were securely strapped at the feet and used the left hand to grasp a fixed bar for support, providing multi-point stabilization of the body. If trunk displacement were the primary source of the observed effects, it would be expected to uniformly attenuate all kinematic measures, particularly in directions where the reaction force exerts a larger torque on the trunk. However, the 45° direction—where trunk perturbation is expected to be minimal according to this account—showed significant changes in movement duration, peak acceleration, and peak speed, while not showing significant timing changes. This dissociation suggests that trunk instability alone cannot account for the observed pattern. Nonetheless, we did not directly measure trunk or shoulder kinematics during the experiment, and we cannot entirely rule out that small trunk displacements contributed to some extent. Future studies would benefit from recording trunk and shoulder motion to more definitively address this factor. Finally, Coriolis and centripetal torques, omitted from our simplified model, might also contribute to movement slowing or direction effect we observed. Previous work has suggested that such effects may be non-negligible in fast planar reaching (Hollerbach & Flash, 1982). To examine their potential contribution, we incorporated Coriolis and centripetal torques terms in our extended model (see Supplementary Notes 2 and Figure 1—figure supplement 4 and 5 for details). These torques were also directionally anisotropic, but their magnitudes were too small to meaningfully affect movement utility or effort computation in our model. Thus, Coriolis and centripetal torques are unlikely to explain the large and systematical effects we observed.

Our results challenge a simple uniform conservative-control account, which posits that movement slowing reflects a generalized adaptation to address stability or safety concerns during spaceflight (Bock, 1998). Arguing against a purely strategic account, taikonauts demonstrated faster reaction times during spaceflight—inconsistent with a generalized slowing strategy. Furthermore, if the sensorimotor system adopted a conservative strategy with pre-planned longer movement durations, the speed profile should maintain symmetry with delayed peak acceleration and speed. Instead, we observed left-skewed speed profiles with earlier peak occurrence for the 90° and 135° directions, where effective mass is greater. This pattern is more consistent with mass-dependent under-actuation than with a uniform strategic adjustment, although we note that the 45° direction did not show a significant timing advance, leaving open the possibility that strategic factors may also play a role in some conditions. This temporal change was more pronounced when the primary submovement was isolated to pinpoint the feedforward control of the reaching since the overall speed profile is affected by the later feedback-based corrections. Previous studies reported inconsistent findings regarding speed profile symmetry, including advanced (Sangals et al., 1999), delayed (Fowler et al., 2008), or unchanged (Berger et al., 1997; Mechtcheriakov et al., 2002) peak speed timing during spaceflight. These discrepancies likely reflect methodological differences, including task variations and technical limitations. For example, two studies used joystick-based pointing tasks without involving limb movement (Fowler et al., 2008; Sangals et al., 1999), which arguably makes them less suitable for examining mass underestimation effects. Mechtcheriakov and colleagues used an outstretched arm task, but their limited sampling rate of 25 Hz may have hindered precise quantification of peak speed, preventing submovement analysis. Additionally, our larger sample size (n = 12) may have enhanced detection of microgravity effects compared to previous studies with typically 3-7 participants (Berger et al., 1997; Bock et al., 2001, 2003, 2010; Fowler et al., 2008; Mechtcheriakov et al., 2002; Papaxanthis et al., 1998; Weber & Stelzer, 2022) or single-participant designs (Heuer et al., 2003; Manzey et al., 1998; Sangals et al., 1999).

If mass underestimation contributes to the observed underactuation, it likely arises through two complementary mechanisms: degraded proprioceptive feedback and misinterpretation of weight-related cues. Microgravity impairs proprioception by reducing both muscle spindle sensitivity (Lackner & DiZio, 1992) and joint receptor responsiveness (Proske & Weber, 2023)—inputs that are crucial for weight perception (Proske & Gandevia, 2012). Furthermore, the sensorimotor system may interpret weight unloading as reduced body mass rather than environmental change (Bock et al., 1996b). Recent studies demonstrate that the sensorimotor system probabilistically infers body-versus-environment causality based on available sensory cues (Berniker & Kording, 2008; Fercho & Baugh, 2014; Kluzik et al., 2008; Kong et al., 2017; Wilke et al., 2013). In normal gravity, consistent mass-weight relationships lead to heavy reliance on weight-related cues for mass estimation (Bock, 1998; Bock et al., 1996b; Sangals et al., 1999). In microgravity, however, reduced weight cues become misleading and may systematically bias mass perception, affecting feedforward control of movement. We note, however, that the same proprioceptive degradation could also affect motor output through other pathways—for instance, by reducing tonic muscle activation or altering spinal reflex gains (Fisk et al., 1993)—independent of any explicit misrepresentation of body mass. Disentangling these mechanisms will require future experiments that can independently manipulate proprioceptive loading and limb inertia.

The persistence of deviant feedforward control and the subsequent within-movement corrections appears intriguing from the theories of human motor learning. Humans demonstrate remarkable ability to adapt to environmental changes in terrestrial studies. For experimentally-applied visual or force perturbations, individuals can recalibrate their sensorimotor control and restore baseline performance within minutes (Krakauer et al., 2019), a process often theorized as forming internal models of the new environment (Kawato, 1999; Wolpert et al., 1995). Relatedly, even when the body mass is perturbed by attaching a weight on the forearm, individuals can fully adapt their reaching movements within tens of trials (Wang & Sainburg, 2004). Previous studies on parabolic flights have also shown that the control of arm movements exhibited rapid adaptation to gravitational changes over the course of several parabolas. (e.g., Gaveau et al., 2016). In contrast, our in-flight sessions were scheduled at least three weeks post-launch—even well beyond the 2-3 weeks window typically considered critical for sensorimotor adaptation in spaceflight (Kanas & Manzey, 2008), but the participants still exhibited un-adapted feedforward control and internal model, suggesting that the underlying sensory bias was not corrected through experience.

We believe this contrast stems from distinct natures of perturbation effects in microgravity when compared to in terrestrial environments. Previous studies on sensorimotor adaptation simulated environmental changes by imposing a perturbation to alter the relationship between motor commands and sensory consequences of motor commands. For reaching adaptation here, visuomotor perturbations changes the spatial mapping between hand motion and its visual representation’s motion (Krakauer et al., 2000; Wolpert et al., 2011); force perturbations change the dynamic mapping between actual force outputs and hand motion (Lackner & Dizio, 1994; Shadmehr & Mussa-Ivaldi, 1994). These mapping changes, though novel, are consistent thus the sensorimotor system is capable of approximating them and adapting accordingly. Theories of sensorimotor adaptation conceptualize the formation of the new mapping as acquiring internal models, which links motor command and efference copy to sensory consequences of motor commands (Flanagan & Wing, 1997; Kurtzer et al., 2008; Wolpert et al., 1998; Wolpert & Miall, 1996). However, the novel environment of microgravity may alter the sensory estimate of motor apparatus (i.e., the body mass here), not merely the sensorimotor mapping. In normal gravity, weight-related sensory cues—mostly proprioceptive feedback—accurately represent body mass. In microgravity, however, these weight-related cues are substantially reduced due to the absence of gravitational pull, providing persistently biased information about body mass. In other words, the microgravity environment failed to provide veridical sensory cues for mass estimation, and reduced gravitational and proprioceptive cues persistently misinform the controller about body mass. Since body mass is a parametric input to the internal models, its bias would affect motor actions even when the internal models are well adapted. Hence, the unique environment of microgravity may reveal an important constraint of the sensorimotor system, i.e., its quick adaptability is restricted to learning a novel sensorimotor mapping, not to persistent sensory bias of bodily property.

The discrete nature of our reaching task may contribute to the persistence of sensory bias. Within-movement corrections indicates that the sensorimotor system learns about sensory bias of mass estimation (Dimitriou et al., 2013), possibly via kinesthetic feedback during movement (Papaxanthis et al., 2005; Proske & Weber, 2023; Weber & Stelzer, 2022). However, movement-related cues only become available after movement initiation. Once a movement ends and the hand returns to rest, microgravity continues to elicit misleading sensory cues that bias mass estimation between trials. This lack of between-trial adaptation parallels findings from a manual interception study aboard the International Space Station, where astronauts persistently initiated movements too early when attempting to intercept an object moving at constant speed (McIntyre et al., 2001). Although astronauts corrected their movements online within each trial, their initial timing error persisted across multiple trials and sessions during a 15-day spaceflight. Hence, discrete motor tasks, such as manual interception and reaching, rely heavily on feedforward control, and are susceptible to sensory biases induced by microgravity. This would also be consistent with the observation that continuous reaching between targets, without stopping, shows quicker adaptation to zero gravity in parabolic flights (Papaxanthis et al., 2005) and does not exhibit prolonged movement duration during spaceflight (Fowler et al., 2008).

What are the potential consequences of the underactuation observed in microgravity beyond the simple reaching movements examined here? The implications may be particularly significant given two key considerations. First, the majority of human movements are not deliberately controlled (Brooks, 1986; Marsden et al., 1976), suggesting that underactuated initial movements may be pervasive in microgravity—especially for actions involving larger body masses such as whole-body movements or object manipulation. Without the controlled conditions and explicit speed requirements of our experimental task, such underactuation might manifest even more prominently in natural movements during spaceflight. Second, our finding of increased reliance on feedback-based corrections points to a potentially costly compensatory mechanism. As feedforward control becomes systematically under-actuated, whether due to mass underestimation, altered neuromuscular properties, or both, the sensorimotor system must depend more heavily on feedback control to achieve adequate performance. This increased reliance on feedback processes may demand additional cognitive resources for action regulation (Bock et al., 2010), potentially contributing to the non-specific stressors that have been shown to impair human performance in space (Tian et al., 2024). The cognitive cost of these compensatory adjustments might be particularly relevant for complex motor tasks or situations requiring divided attention during spaceflight operations.

The preservation of speed-accuracy trade-offs in our study extends previous findings from Fitts’ task experiments performed during spaceflight (Fowler et al., 2008). Both the traditional speed-accuracy trade-off and action planning-dependent trade-off remained intact in microgravity (Figure 2—figure supplement 3 and 4), suggesting preserved fundamental motor control capacity. Motor performance during spaceflight typically degrades in tasks with high cognitive demands (Eddy et al., 1998; Manzey et al., 1998; Strangman et al., 2014), particularly those requiring sustained attention (Tian et al., 2024). Our findings suggest that while microgravity-induced underactuation affects movement execution, the underlying motor control capabilities remain robust in microgravity when cognitive demands are modest.

Our study presents four methodological limitations that warrant consideration. First, despite matching for age and gender, our ground controls exhibited faster movements than the taikonauts. This systematic difference, reflected in higher peak acceleration and speed, may stem from differing experience with the touchpad device used for movement recording. While taikonauts used touchpads regularly, 9 of 12 control participants had minimal experience, potentially leading to compensatory faster movements to avoid overtime errors. Second, the temporal evolution of mass underestimation effects remains unclear. Our earliest measurements occurred three weeks post-launch, though parabolic flight studies suggest these effects may emerge within hours of microgravity exposure (Crevecoeur et al., 2014; Papaxanthis et al., 2005). Additionally, incomplete post-flight recovery in some kinematic measures suggests either a mass overestimation due to exaggerated weight-related cues or neuromuscular deficiency typically observed shortly after returning to earth (Tays et al., 2021). Third, while our hand-reaching task enabled us to use well-established analytical frameworks for studying motor control aspects of movements, the generalizability of mass underestimation effects to other movements, particularly whole-body actions, remains to be determined. Future research should address these limitations by implementing matched device training protocols, adding early-and post-flight measurements, and extending investigations to diverse motor tasks. We also notice that the microgravity effect was less consistent for the 45° reaching direction. This direction engages more single-joint, elbow-dominant reaches, whereas the 90° and 135° targets require greater multi-joint (shoulder + elbow) coordination. Consistent with this, an exploratory analysis of hand-path curvature (cumulative curvature; positive = counterclockwise) showed larger curvature at 45° (6.484° ± 0.841°) than at 90° (1.539° ± 0.462°) or 135° (2.819° ± 0.538°). The significantly larger curvature in the 45° condition suggests that these movements deviate more from a straight-line path, supporting a more elbow-dominant movement in this direction. Importantly, this curvature pattern was present in both the pre-flight and in-flight phases, indicating that it is a general movement characteristic rather than a microgravity-induced effect. We postulate that single-joint movements, more so than we considered in our ideal two-link model, made effective mass rather small in the 45° direction, hence the mass underestimation and underactuation in feedforward planning had smaller effect, compared to model simulations, in this direction. Fourth, our interpretation of the observed underactuation relies on a simplified optimal control model that treats muscles as ideal torque generators. This model does not capture several neurophysiological factors that may be altered in microgravity, including changes in tonic muscle activation, spinal reflex modulation, and descending vestibular influences on motor neuron excitability (Fisk et al., 1993), nor potential alterations in the damping and natural frequency of the limb’s mechanical response. However, the available evidence suggests that upper limb muscle capacity is relatively preserved in microgravity: a systematic review found that upper limb maximal voluntary contraction remained mostly unchanged during unloading periods of up to 45 days, with upper limb muscles declining substantially more slowly than lower limb and trunk muscles (Winnard et al., 2019; Bosutti et al., 2025). While the direction-dependent pattern of our results is most parsimoniously explained by a misrepresentation of inertial properties, we cannot exclude that some portion of the observed underactuation arises from these other neuromuscular factors. Distinguishing between mass underestimation and other sources of underactuation would benefit from future studies that combine detailed musculoskeletal modeling with direct measurements of muscle activation, joint impedance, and trunk kinematics during spaceflight.

In conclusion, our study provides converging evidence that movement slowing in microgravity is associated with systematic underactuation of initial movements, a pattern most consistent with the underestimation of body mass hypothesis. By leveraging the direction-dependent variation in limb effective mass, we showed that reaching movements in microgravity exhibit kinematic signatures—reduced and earlier peak speed/acceleration—that are broadly consistent with the predictions of the mass underestimation hypothesis. Although other neurophysiological factors such as altered muscle tone and vestibular modulation may also contribute, these effects are not readily explained by a uniform strategic adjustment or general neuromuscular change alone. The persistence of these effects throughout long-duration spaceflight, coupled with increased feedback-based corrections, suggests how altered sensory inputs can impact both predictive and reactive components of motor control in ways that resist sensorimotor adaptation. While basic motor control capabilities remain intact, as evidenced by preserved speed-accuracy trade-offs, the consistent underactuation of movements may have significant implications for the theorization of sensorimotor learning and for practical considerations of motor performance during space operations. These findings not only advance our understanding of how the sensorimotor system adapts to novel gravitational environments but also highlight fundamental principles of sensorimotor integration, particularly the central role of gravitational cues in shaping our perception of body dynamics.

## Methods

### Participants

Twelve taikonauts (2 females, 10 males; mean age 49.5 ± 6.5 years) from the first four missions of the China Space Station (CSS) served as the experimental group (Table S1). The control group consisted of 12 right-handed residents from Beijing (2 females, 10 males; mean age 49.9 ± 6.5 years). Control group received monetary compensation. All participants were provided with written informed consent forms, approved by the Ethics Committee of the China Astronaut Research and Training Center and the Institutional Review Board of Peking University. The experimental group was exposed to microgravity for 92–187 days during manned missions in China’s Shenzhou Program. Due to operational constraints, the number of test sessions varied among taikonauts: one completed 4 sessions, five completed 6, and six completed 7 (Table S1). All control sessions were conducted at Peking University.

### Task and experimental design

The task was conducted using a tablet (Surface Pro 6, Microsoft, Redmond, Washington) with 100Hz sampling frequency and noise-canceling earphones (FiiO FA7, FiiO Electronics Technology, Guangzhou, China). During in-flight sessions, participants wore earphones and assumed a neutral posture with feet secured by foot straps in front of a foldable tabletop, positioned 0.9 meters above the cabin floor. They held the tabletop edge with their left hand, allowing their right hand to move freely on the tablet (Figure 7A). The foot straps and left-hand grip provided body stabilization. The tablet, attached to the tabletop with Velcro, was placed ~35 cm in front of participants and 30 cm below chest level (Figure 7A). At the start of each trial, an orange circle (0.5 cm radius) appeared at the bottom of the screen, signaling participants to move their right index finger to this start circle. After a random delay (500–1000 ms), a target appeared at one of three possible locations, 12 cm from the start circle at 45°, 90°, or 135° counterclockwise from the horizontal axis (Figure 7A). Participants were instructed to reach the target quickly and accurately, pausing briefly before returning to the start. If the time from target appearance to movement end (finger speed <10 cm/s) exceeded 650 ms, a “too slow” message prompted faster movement. In half of the trials, a 50 ms “beep” sound was played at target appearance. Each session included 120 trials, with 3 target directions and 2 beep conditions balanced and randomized (Figure 7B). The x-y position of the index finger was recorded continuously at 100 Hz.

**Figure 7.**
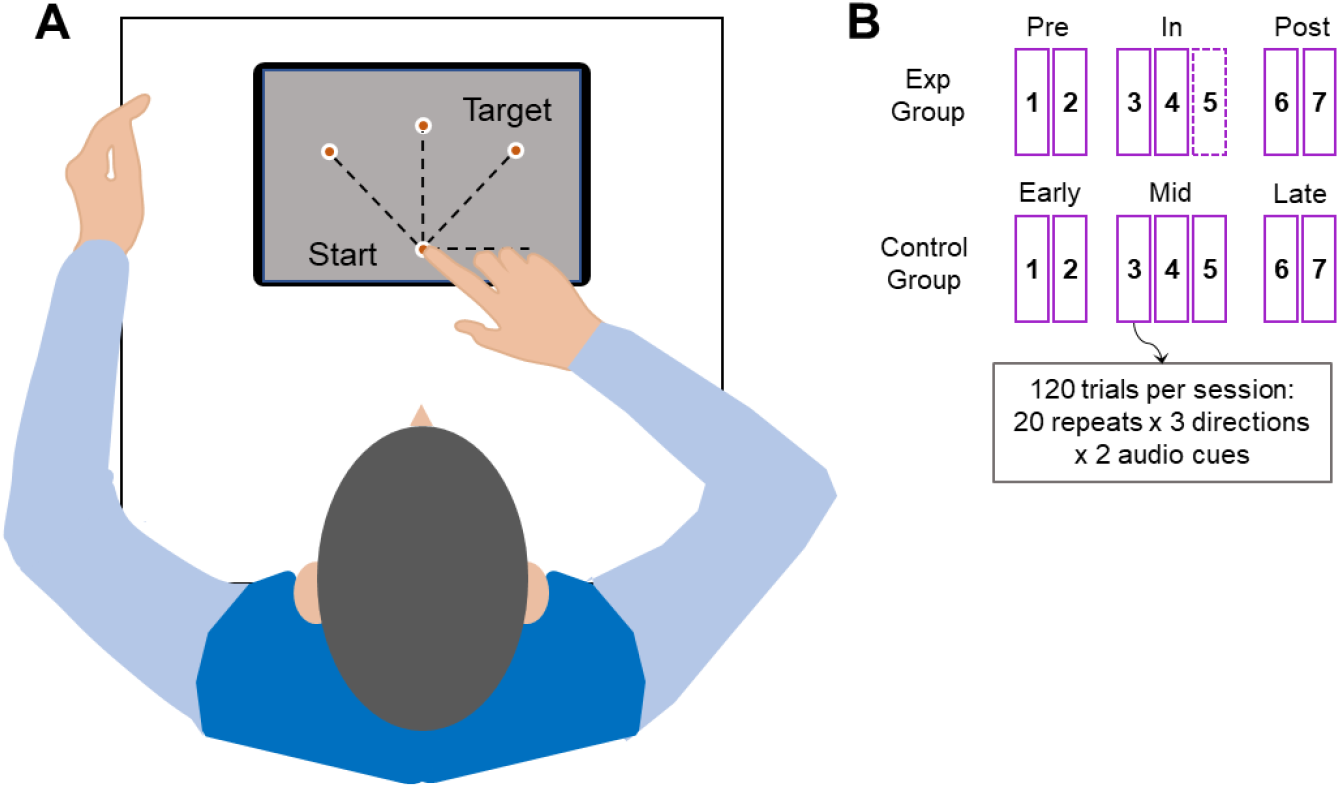
Experimental Setup and Design. **A)** Top-down view of a participant performing the reaching task with the right hand on a tablet. The start position and all possible target locations are shown as orange dots on the tablet screen. **B)** Experimental design. Both groups completed 4–7 sessions, with each session consisting of 120 trials. Some taikonauts missed 1 or 2 in-flight sessions.

All experiments aboard the China Space Station were monitored in real time by experimenters at the Beijing Control Center. Task instructions were displayed on the opening screen of the data-acquisition application, and participants were given ample time to read them. The same team of experimenters administered all pre-, in-, and post-flight sessions using identical instructions. Consistent with standard practice, the astronauts served as both participants and on-orbit experimenters and were extensively trained for this role on the ground. Multiple pre-flight sessions were conducted to familiarize them with the task. These safeguards were implemented to ensure high-quality data.

The control group was included to assess a potential confounding effect of repeated measurements. The interval between successive test sessions was 6–9 days, which was shorter than those of the taikonauts (1–2 months) to conservatively evaluate potential practice effects. Despite the shorter intervals, no significant differences were observed across sessions for most measures (except reaction time), confirming minimal practice effects and providing a robust baseline for evaluating microgravity’s impact on the taikonauts. All ground tests, for both groups, were conducted using identical devices and software. The layout of the table and tablet was kept consistent across sessions. The only difference was that, in ground tests, participants sat in a chair approximately 90 cm above the ground without foot straps and did not need to hold the edge of the tabletop with their left hand for stability.

### Model simulation

We simulated the peak speed, peak acceleration, and their corresponding times for reaching movements in different directions. We employed the classical two-joint arm model to demonstrate the effect of biomechanics (Li & Todorov, 2004; Figure 1B). This was combined with a movement utility model to estimate planned movement time (Shadmehr et al., 2016; Figure S6A and S5B) and a forward optimal controller to simulate reaching movements (Todorov, 2005; Figure 1C and F, Figure S6C-D). Our simulation is briefly outlined below in steps.

We used a two-joint planar arm model to estimate the effective mass of the hand moving in different directions (Figure 1B). The parameters of the upper arm (length *l*_1_, mass *m*_1_, center of mass 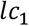, inertia *I*_1_) and the lower arm (length *l*_2_, mass *m*_2_, center of mass 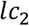, inertia *I*_2_) are based on values from Shadmehr et al., 2016:

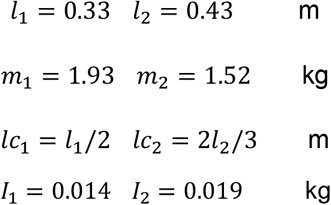

The elbow and shoulder angles are denoted by *θ*_*e*_ and *θ*_*s*_, respectively. The initial arm configuration is set by *θ*_*s*_ = 0.785 rad, and *θ*_*e*_ = 1.571 rad, following the experimental conditions from Gordon et al., 1982. With a particular joint angle configuration, the end-point hand position is:

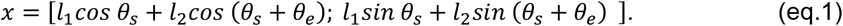

The arm’s inertia matrix is:

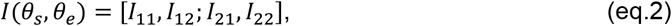

where:

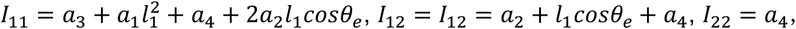

with:

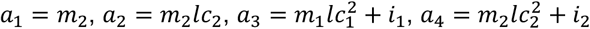

The joint torques at the shoulder and elbow are represented as:

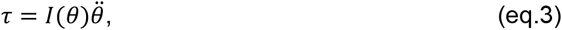

and the forces at the hand are:

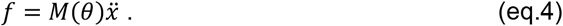

We use the Jacobian matrix 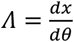 to relate force and torque, based on virtual work principle:

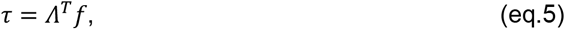

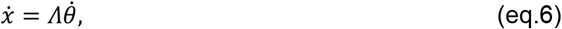

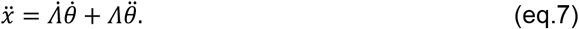

Put S3 into S5:

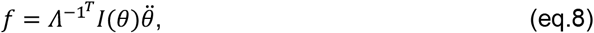

then combine S7 and S8:

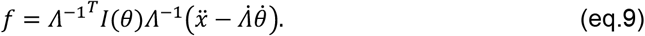

Since the hand speed at the beginning of the movement is zero, according to S4 and S9, the mass matrix of hand *M*(*θ*) is:

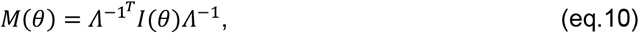

The mass matrix *M*(*θ*) is a 2 x 2 matrix, and the effective mass *m*(*θ*) is determined by the length of the force vector when *M*(*θ*) is subjected to a unit acceleration.

In Figure S6A, the solid curve illustrates the effective mass across various directions under normal gravity, while the dashed curve represents the effective mass with a hypothetical 30% 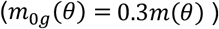 mass underestimation in microgravity. The colored lines indicate the effective mass amplitudes for the three target directions used in our experiment: 45°, 90°, and 135°.

#### Movement time determined by optimal utility

Following Shadmehr et al., 2016, the utility of an reaching action is given by:

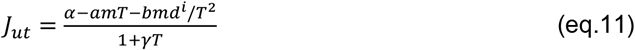

where *α* is the reward, m is mass, d is the distance to be moved, T is the planned duration of the movement, and a, b and i are scaling factors of the effort cost. *γ* represents a temporal discounting factor. The optimal movement time (*T*_*opt*_) is the time that maximizes the utility *J*_*ut*_ :

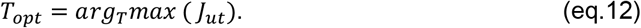

The parameter values used in the simulation of Figure S6B were taken from Shadmehr 2016: *α* = 1000, *a* = 15, *b* = 100, *i* = 1.1, γ = 1. The movement distance *d* is set as 0.12 m, the target distance in the experiment. Figure 1—figure supplement 1B illustrates how the utility curves change with planned movement time in three experimental directions (45°, 90°, and 135°) under normal gravity, with colored dots indicating the optimal movement durations.

*Movement kinematics simulated by optimal control theory*. After setting the movement time *T*_*opt*_, the position, speed and acceleration over time can be simulated based on an optimal control model (Todorov, 2005). The arm’s control operates as a second-order low-pass filter:

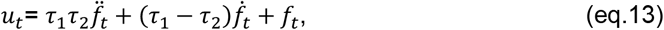

with the time constants τ_1_ = τ_2_ = 40*ms*.

The system state vector is 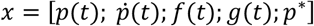, where p is position of hand, f is force acting on the hand, and g is an auxiliary state variable, p* is the target position. The control law is given by:

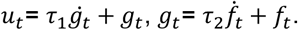

The system dynamics are:

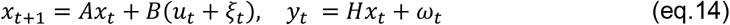

With the sensory feedback carries information about position, velocity and force:

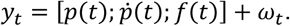

ξ_*t*_ is signal-dependent noise 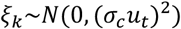. ω_*t*_ is sensory noise ω_*k*_~*N*(0, Ω^ω^), where

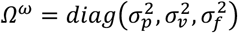

We define the sampling interval Δ*t*, and the number of steps is *N* = *T*_*opt*_/Δ*t*, the discrete-time dynamics of the above system are:

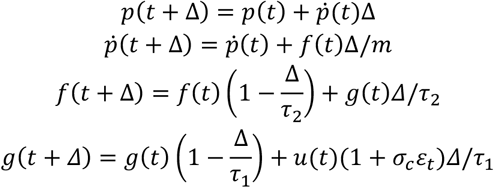

Which is transformed into the matrices as:

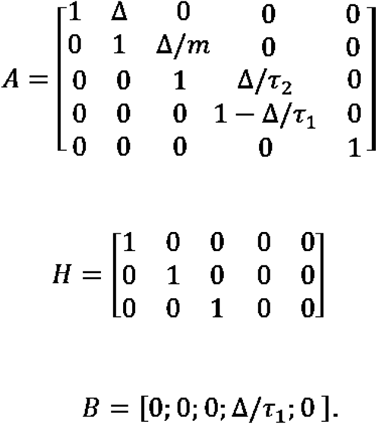

The controller seeks to minimize the cost function J by optimizing the command u:

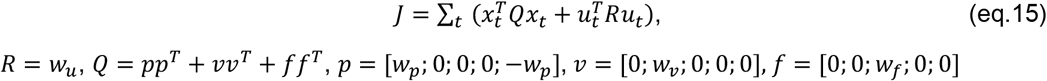

Here, we used the penalty values from Todorov (2005) for each state and control variable: 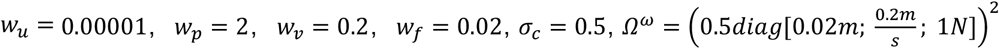. We first estimated the time-varying gains {*L*_*k*_, *K*_*k*_} correspond to the feedforward mapping and the feedback correction gain, respectively. The control law can be expressed as: 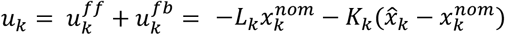, where *u*_*k*_ is the control input, 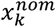 is the nominal planned state, 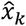 is the estimated state, *L*_*k*_ is the feedforward (nominal) control associated with the planned trajectory, and *K*_*k*_ is the time-varying feedback gain that corrects deviations from the plan. To define the motor plan for comparison with behavior, we then simulate the deterministic open-loop trajectory by turning off noise and disabling feedback corrections, i.e., 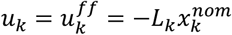. We then obtain the peak speed, peak acceleration, and their respective times.

In summary, we apply the movement utility theory to calculate the optimal movement time for a simplified two-joint arm moving in a 2D plane. This movement time is primarily determined by the effective mass and temporal discounting of reward. Next, we employ the optimal control theory to model the kinematics of hand movement, including its position, speed, and acceleration. This approach also enables us to estimate the peak speed and acceleration, as well as their respective times, for reaches performed toward different target directions. To simulate the effect of microgravity, we introduce a mass underestimation factor to account for the kinematic changes associated with this perceptual error.

## Data analysis

*Kinematic analysis*. A Butterworth low-pass filter with a 20 Hz cutoff frequency was applied to the positional data. Movement distance at each time point was computed as the Euclidean distance between the finger’s current position and the center of the start circle. Speed was estimated by performing linear regression over a moving window of five consecutive positional data points, and acceleration was calculated similarly from the speed data. Movement onset was defined as the first time point at which acceleration exceeded 50 cm/s2, while movement offset was the first time point at which acceleration fell below 50 cm/s2. Reaction time was defined as the interval between target onset and movement onset, and movement duration as the interval between movement onset and offset. Movement endpoint error was computed as the Euclidean distance between the target and the finger’s position at movement offset. Peak speed and peak acceleration were identified as the maximum speed and acceleration occurring between movement onset and offset, respectively. The times of peak speed and peak acceleration were measured as their respective time intervals from movement onset, and their relative times were calculated by dividing each by the movement duration.

*Submovement extraction*. To separate the primary submovement from the secondary corrective submovement, the speed profile of each reaching movement was decomposed into a sum of submovements. Following the optimization algorithm described by Rohrer and Hogan (2006), the speed profile was fitted as a sum of support-bounded lognormal (LGNB) curves. Each LGNB curve is defined as:

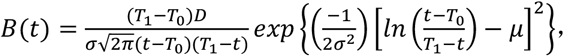

where *T*_*0*_ and *T*_*1*_ represent the start and end times of the movement, D is a scaling parameter, and the parameters μ and σ determine the skewness and kurtosis of the underlying lognormal function. These five parameters enable each LGNB submovement to vary across a wide range of possible shapes.

In theory, the number of submovements could be treated as a free parameter. However, to avoid overfitting, we limited each movement to two submovements, given the brevity of the movement (average duration approx. 350 ms). For each trial, we initially fit the data using both one and two LGNB curves and then determined the optimal number of submovements using the “greedy” method from the previous study (Rohrer & Hogan, 2006). Briefly, the fitting error for each movement with one and two LGNB curves was calculated as:

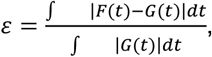

where G(t) is the actual movement speed profile, and F(t) is the fitted speed profile (the sum of the submovements). If the fitting error for a single submovement was below a 2% threshold, or if adding a second submovement reduced the fitting error by less than 2%, a single submovement was considered adequate. Otherwise, two submovements were included. Trials with large fitting errors (>10%) were discarded, accounting for 3.60% and 4.83% of trials in the experimental and control groups, respectively. The percentage of trials with two submovements was calculated for each movement direction and phase. The time difference between the peaks of the two submovements was also computed, and set to zero if only one submovement was detected. Data fitting was performed using Matlab’s *fmincon* function (MathWorks, Natick, MA, USA), and the robustness of fitting was confirmed by testing 20 random sets of initial parameters.

*Statistical analysis*. First, invalid trials were excluded from further analysis, with three possible causes: early initiation (RT < 150 ms, indicating guessing of the target direction), affecting 0.74% and 0.70% of trials in the experimental and control groups, respectively; late initiation (RT > 400 ms, indicating inattention), affecting 1.13% and 0%; and measurement failures, affecting 1.16% and 0.70%. The proportion of invalid trials did not differ significantly across testing phases (Friedman’s tests, all p > 0.1).

Most dependent measures—including movement duration, peak acceleration, peak speed, time to peak acceleration/speed, relative time to peak acceleration/speed, the percentage of submovements, and inter-peak intervals—were analyzed using two-way repeated-measures ANOVAs with a 3 (phase) × 3 (movement direction) design. Reaction time was analyzed using three-way repeated-measures ANOVAs with a 3 (phase) × 3 (movement direction) × 2 (beep/no beep) design. Greenhouse-Geisser corrections were applied when sphericity was violated (Kirk, 1968). Normality was assessed with Shapiro-Wilk tests, and if more than 10% of tests for a dependent variable violated normality, the Aligned Rank Transform (ART) procedure was applied before conducting repeated-measures ANOVA. Tukey’s HSD tests were used for pairwise comparisons.

To examine the relationship between submovement and movement duration, a Linear Mixed Model (LMM) was used to analyze the association between changes in the inter-peak interval of submovements (ΔIPI) and changes in movement duration (ΔMD). The dependent variable *y*_Δ*MD*_ represents the mean change in movement duration across two adjacent phases (pre-to-in and post-to-in transitions) for each target direction. Target direction (45°, 90°, 135°) and phase transitions (pre-to-in and post-to-in) were treated as within-subject factors. The predictors included three fixed factors: ΔIPI, phase transition (*x*_*pha*_), and direction (*x*_*dir*_). An intercept term (*u*_*par*_) was included as a random factor to account for individual differences in movement duration. The resulting LMM is as follows:

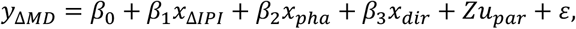

where β_0_ denotes the overall intercept and ε denotes a Gaussian residual error.

All statistical analyses were performed using MATLAB’s Statistics and Machine Learning Toolbox. The significance level was set at α = 0.05, and all tests were two-tailed unless otherwise specified.

For the directional dependent effects (Figure 6), we applied repeated measures correlation to quantify within-subject associations (Bakdash & Marusich, 2017). Each participant contributed three repeated measurements under different directions. The predictor variable was identical across conditions, and the outcome variables were the measured responses of interest. Repeated measures correlations were computed separately for each of the four metrics, controlling for between-subject differences by including subject-specific intercepts and estimating a single common slope across participants.

## Acknowledgements

We would like to thank Mr. Shaolin Zeng for his help with data acquisition of the control group.

## Authors contributions

Conceptualization: KW, YT, ZZ; Data curation: YT, ZZ; Formal analysis: ZZ; Funding acquisition: KW, CW, YT, ZZ; Methodology: KW, ZZ; Software: YT, CW, CJ, BW, HY, RZ; Supervision: KW; Visualization: ZZ; Writing: KW, ZZ.

## Data and materials availability

Control group data are deposited in OSF (https://osf.io/3ac5b/overview?view_only=4b93189789e34c178283e8b99550266e). Analysis code is available on GitHub (https://github.com/ZhaoranZhang/Aiming_data_analysis.git). Raw astronaut data are restricted under China’s manned space program confidentiality regulations and cannot be shared publicly. Requests for access for scientific research purposes may be directed to the corresponding author, subject to approval by the China Astronaut Research and Training Center.

## Supplemental Materials

**Figure 1—figure supplement 1.**
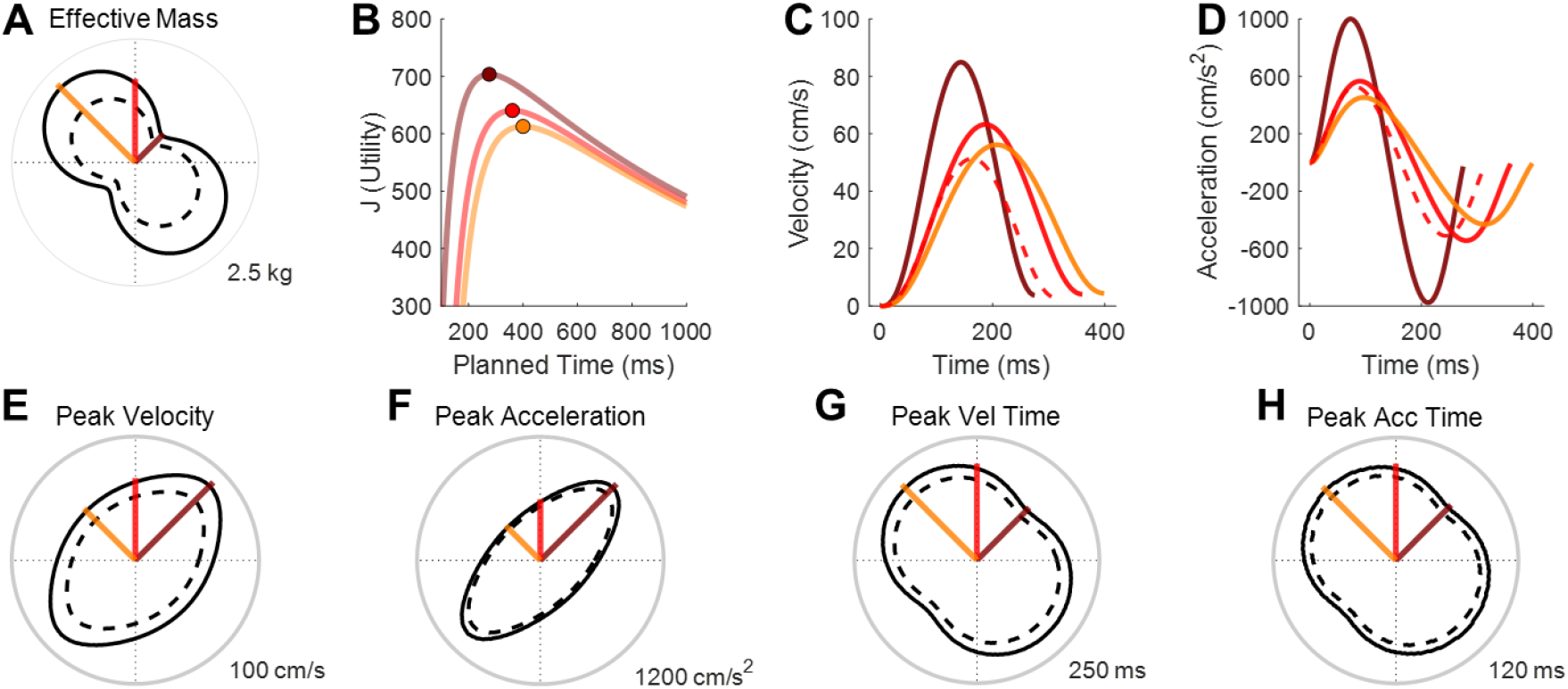
Simulation details for the arm model. **A)** Effective mass across different movement directions, with three colored lines indicating the specific directions used in the experiment. The solid-line contour denotes the effective mass simulated by default parameters under normal gravity (see Supplemental Text 1), and the dash-line contour denotes a hypothetical 30% underestimation of the effective mass in microgravity **B)** Utility function values for movements in each of the three directions. **C-D)** Speed and acceleration profiles for movements in these directions. **E-H)** Simulated values of peak speed, peak acceleration, peak speed timing, and peak acceleration timing across all movement directions. The solid-line contours denote simulated values by default parameters, and the dash-line contours denotes the values with a 30% mass underestimation.

**Figure 1—figure supplement 2.**
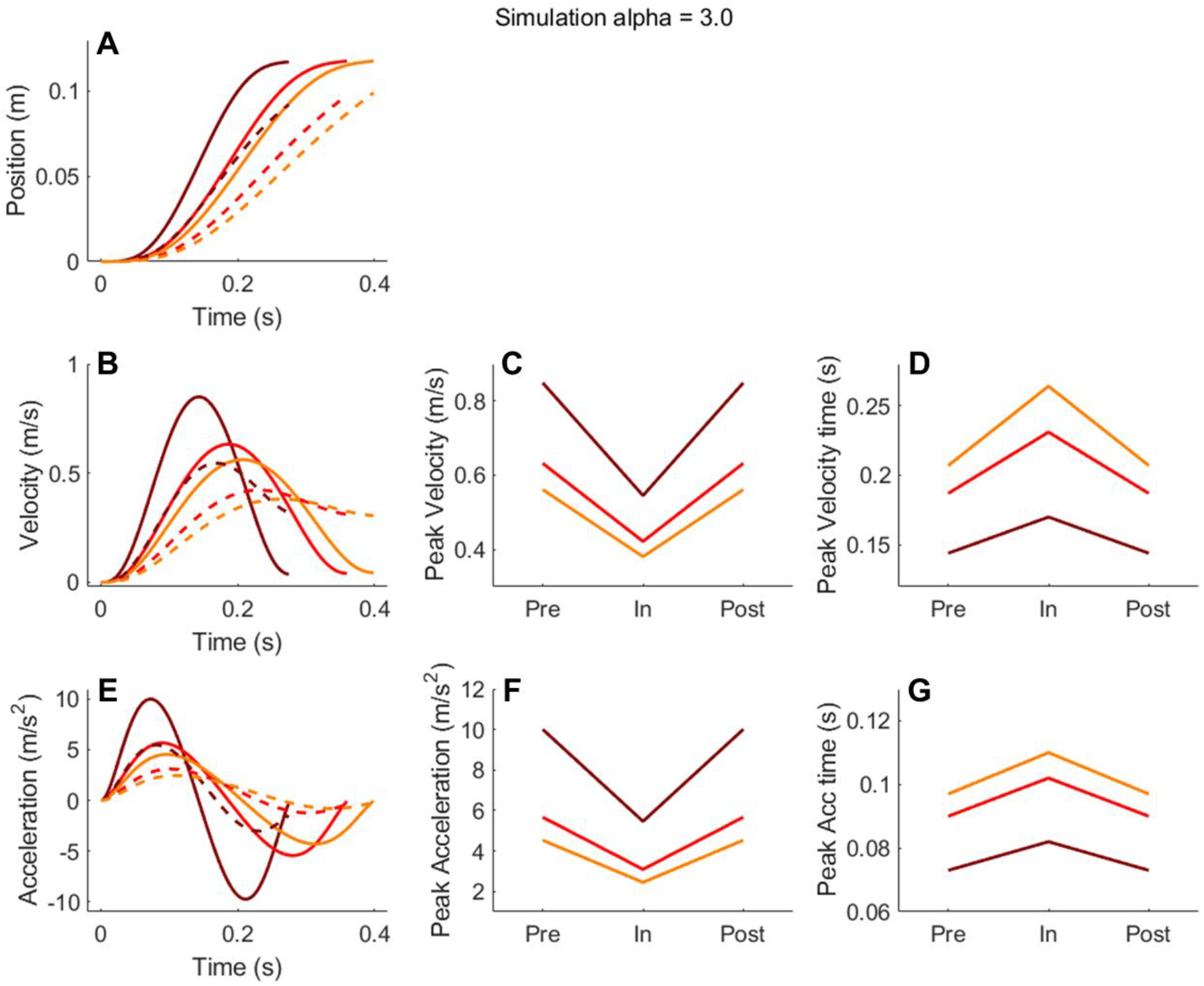
Simulation using an altered cost function with α = 3.0. Panels A, B, and E show simulated position, velocity, and acceleration profiles, respectively, for three movement directions. Solid lines correspond to pre- and post-exposure conditions, while dashed lines represent the in-flight condition. Panels C and D display the peak velocity and its timing across the three phases (Pre, In, Post), and Panels F and G show the corresponding peak acceleration and its timing.

**Figure 1—figure supplement 3.**
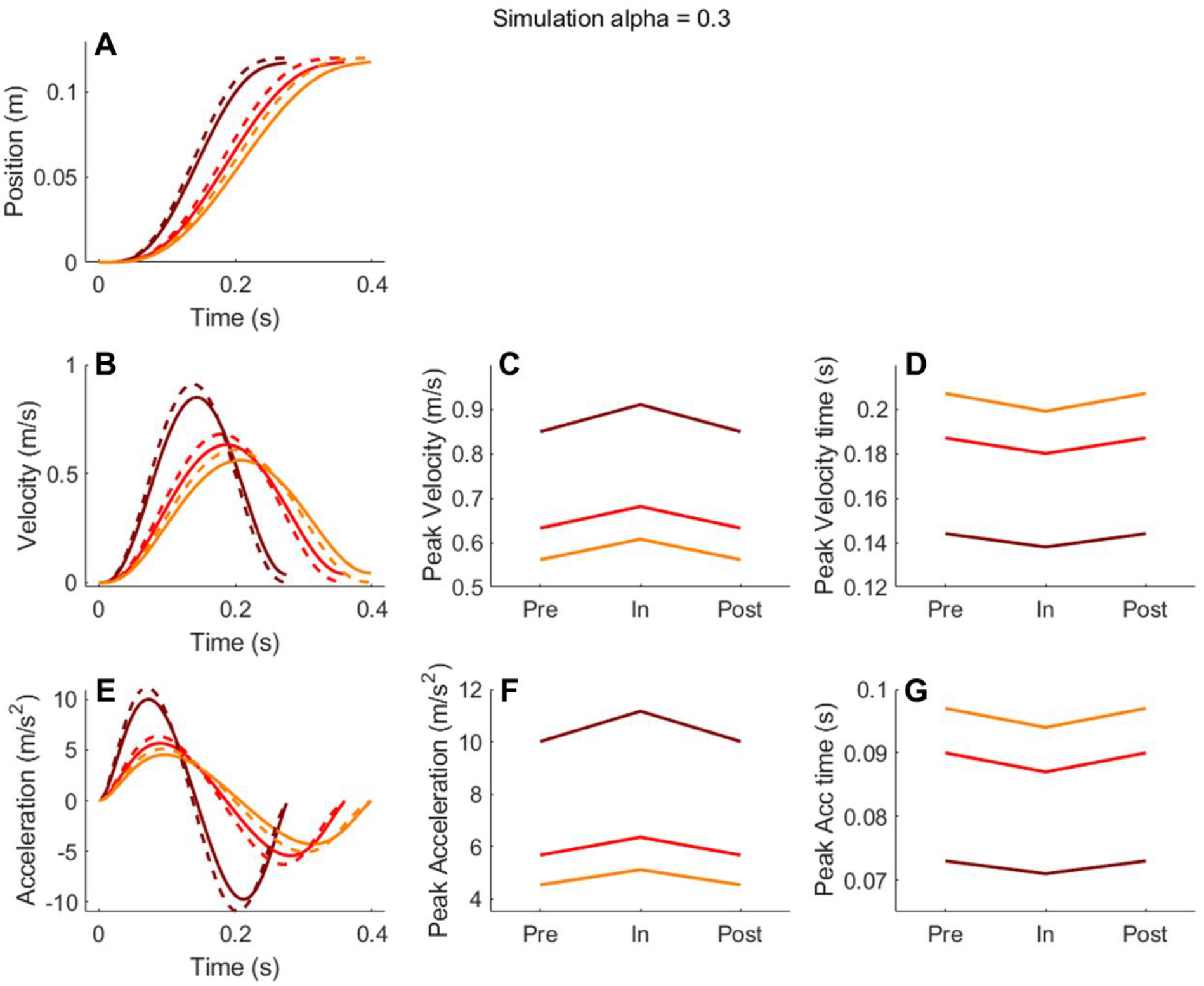
Simulation results under altered cost function with α = 0.3. Same as Figure 1—figure supplement 2.

**Figure 1—figure supplement 4.**
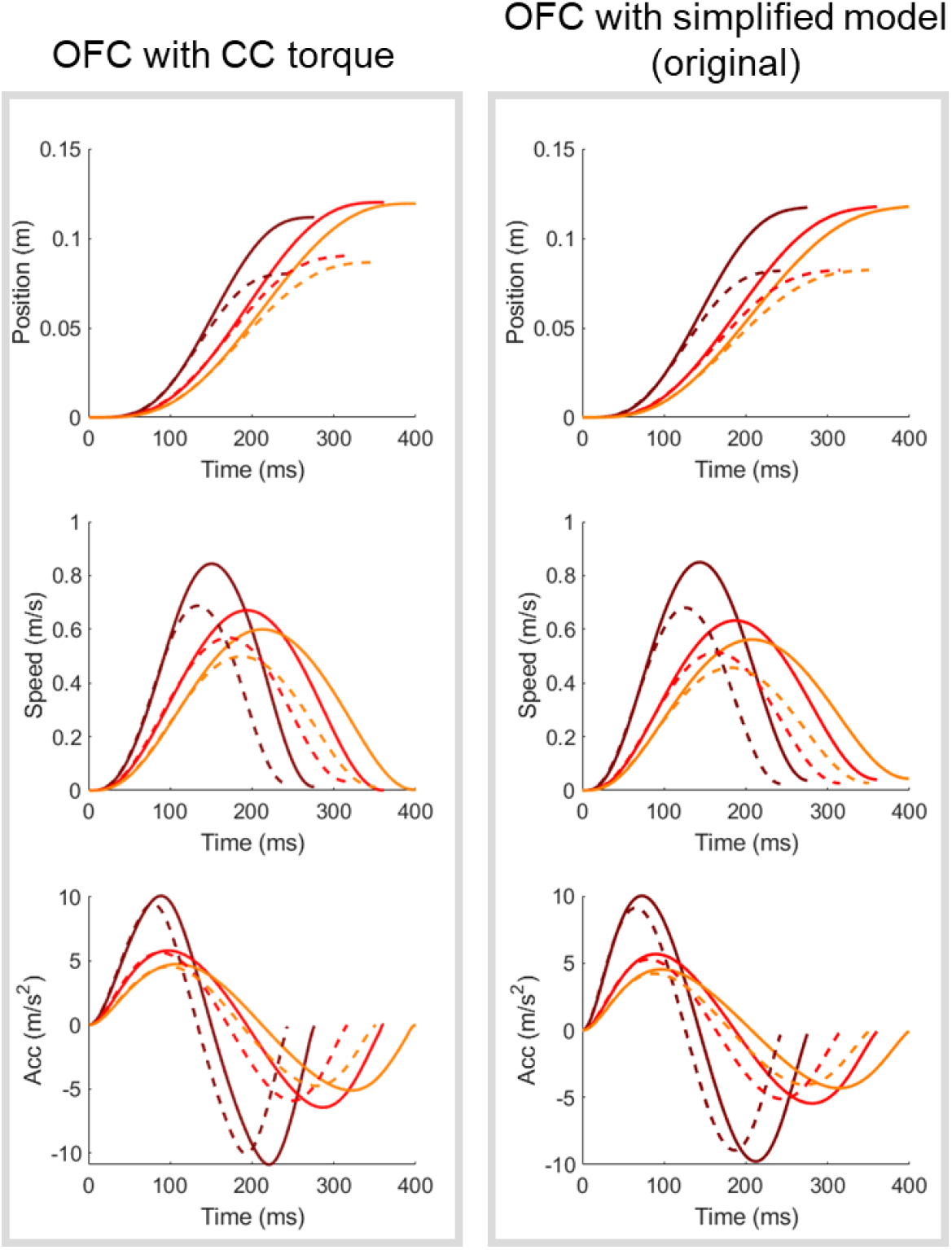
Comparison between simulation results from the model with CC torque and the simplified model. The left panels show results from the model with CC torque, and the right panels show those from the simplified model. The top row displays the position profiles, and the middle and bottom row shows the corresponding speed and acceleration profiles. The three colors represent three movement directions (dark red: 45°, red: 90°, yellow: 135°). Dashed lines indicate the simulated trajectories under microgravity, assuming mass underestimation. For the full model, we used a 2-link arm model that includes both inertia and Coriolis and centripetal torques (Supplementary Note 2).

**Figure 1—figure supplement 5.**
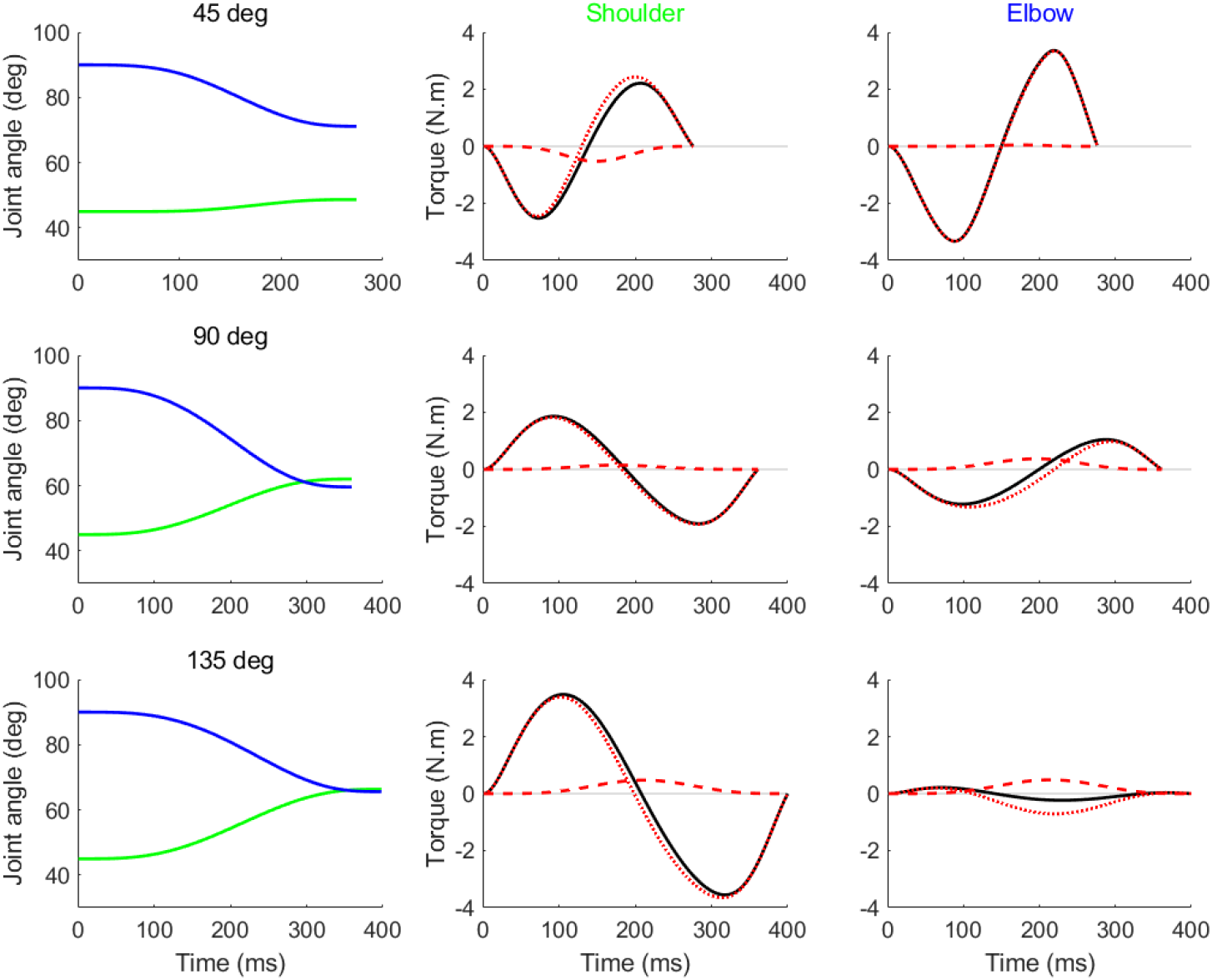
Joint angles and joint torques of shoulder and elbow with simulated trajectories towards three different directions. **A)** Shoulder (green) and elbow (blue) angles change with time for the 45° movement direction. **B)** Components of joint interaction torques at the shoulder. Solid line: net torque at the shoulder; dotted line: shoulder inertia torque; dashed line: shoulder Coriolis and centripetal torque. **C)** Same plot as B for the elbow joint. **D–F)** Same as A–C for the 90° movement direction. **G–I)** Same as A–C for the 135° movement direction.

**Figure 2—figure supplement 1.**
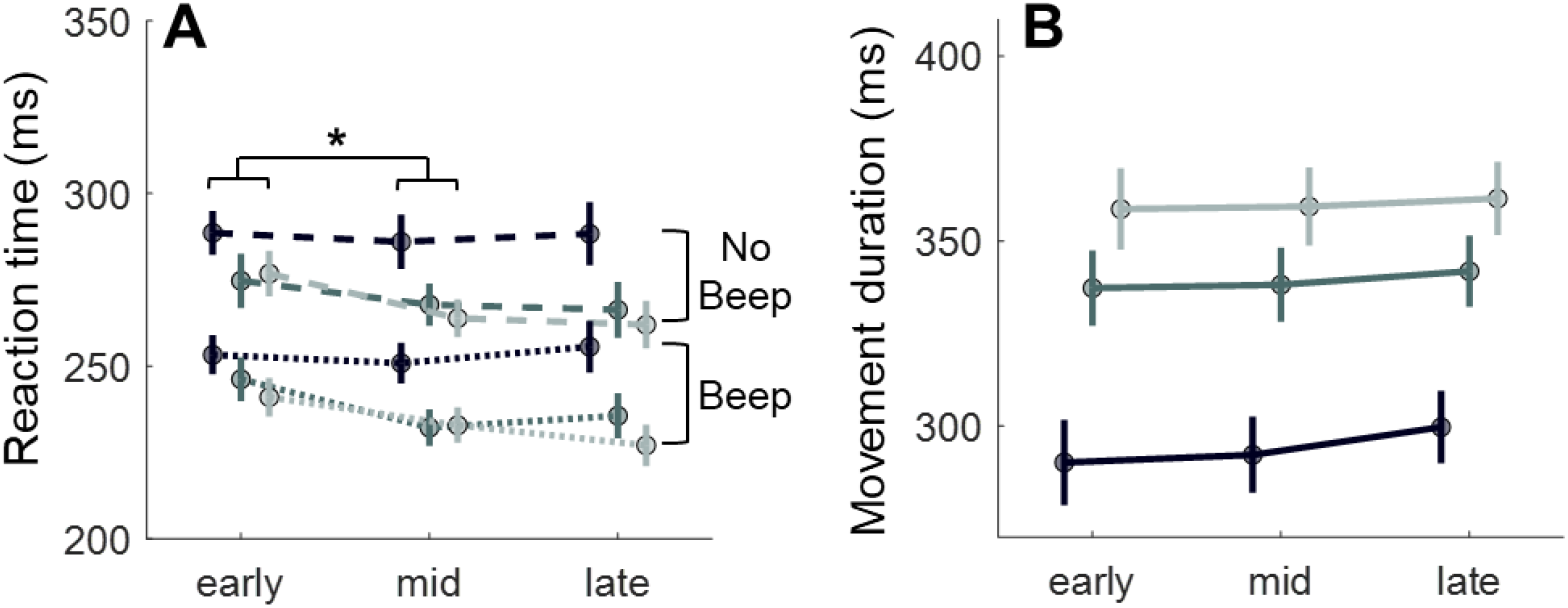
Reaction time and movement duration in the control group. The control group exhibited no phase effect for the movement duration, and a slightly decrease in reaction time. However, similar effects were also observed in the experimental group. Each measure was analyzed using a 3 (direction) × 3 (phase) two-way repeated-measures ANOVA. **A)** Reaction Time: Significant main effects of direction (F(2,22) = 21.946, *p* < 0.001, partial η2 = 0.666) and phase (F(2,22) = 4.281, *p* = 0.039, partial η2 = 0.280) were found, along with a significant direction-phase interaction (F(2,22) = 7.940, *p* = 0.001, partial η2 = 0.419). RT was significantly faster in the Middle phase than in the Early phase (*p* = 0.032, D = 0.589). **B)** Movement Duration: Significant main effect of direction (F(2,22) = 99.874, *p* < 0.001, partial η2 = 0.901), with all pairwise comparisons significant at *p* < 0.001. Neither main effect of phase (F(2,22) = 0.995, *p* = 0.360) nor interaction effect (F(4,44) = 1.412, *p* = 0.114) was significant.

**Figure 2—figure supplement 2.**
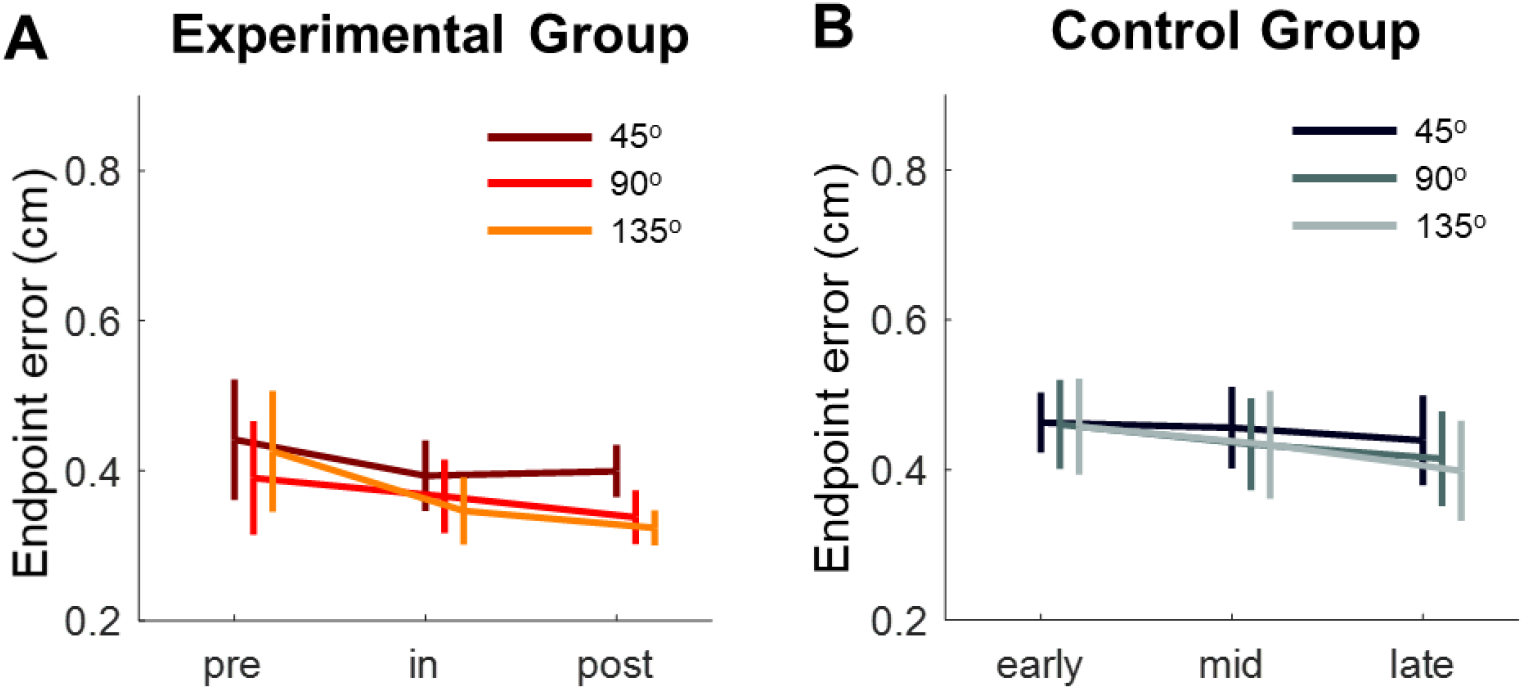
Movement accuracy quantified by endpoint error. The average endpoint error for each target was analyzed across experimental phases. Both the experimental group **A)** and control group **B)** performed the task with high accuracy despite the stringent timing requirements. The presence of a beep had no effect on endpoint error in either group (experimental group: main effect, F(1,11) = 1.4461, *p* = 0.254; all interactions, *p* > 0.2; control group: main effect, F(1,11) = 0.717, *p* = 0.415; all interactions, *p* > 0.25). Therefore, data from the two beep conditions were pooled, and a two-way repeated-measures ANOVA was conducted for each group. In the experimental group, endpoint error showed a slight decrease across phases, but the main effect of phase was not statistically significant (F(2,22) = 1.768, *p* = 0.210), nor was the interaction effect (F(2,22) = 2.386, *p* = 0.091). However, there was a significant main effect of direction (F(2,22) = 16.020, *p* < 0.001, partial η2 = 0.593), with post-hoc comparisons revealing that the 45° direction had significantly larger endpoint errors than the 90° and 135° directions (*p* = 0.002, partial η2 = 0.291; *p* = 0.001, partial η2 = 0.286, respectively). In the control group, no significant effects of phase or direction were observed, and there was no significant interaction (all *p* > 0.23).

**Figure 2—figure supplement 3.**
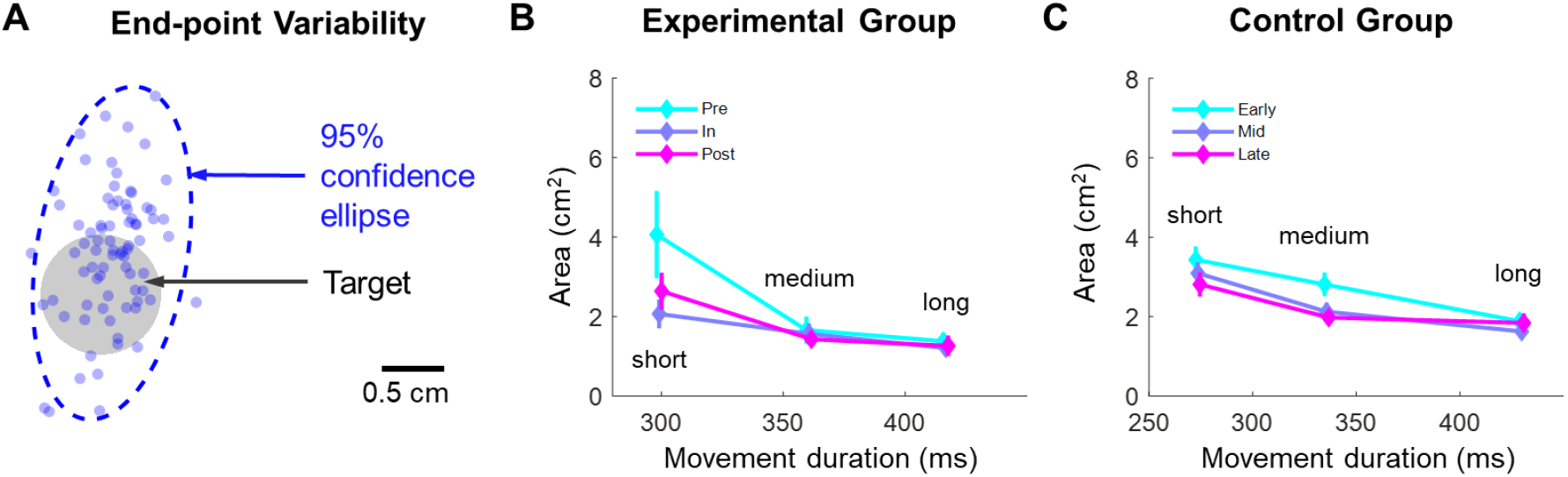
Speed-accuracy trade-off as indicated by the relationship between movement duration (MD) and endpoint dispersion, following the classical Fitts’ law. **A)** Endpoint dispersion was estimated by constructing a 95% confidence ellipse encompassing the endpoints of all trials. Regardless of movement direction, phase, and beep condition, trials were divided into three equal-sized bins based on duration, labeled as short, medium, and long. For each participant at each phase, trials within each bin were pooled to estimate the ellipse of endpoints. **B-C)** A positive association between endpoint area and MD was observed, indicating a speed-accuracy trade-off. Both groups exhibited a dependency of the endpoint area on MD. For the experimental group, a 3 (phase) × 3 (MD) two-way repeated-measures ANOVA on endpoint area showed a significant main effect of MD (F(2,22) = 14.125, *p* = 0.003, partial η2 = 0.407), with only a marginal phase effect and no interaction (phase: F(2,22) = 3.199, *p* = 0.087; interaction: F(4,44) = 2.106, *p* = 0.163). Pairwise comparisons indicated that short, medium, and long MD conditions significantly differed from each other (short vs. medium: *p* = 0.018, D = 1.030; medium vs. long: *p* = 0.007, D = 0.202; short vs. long: *p* = 0.004, D = 1.232). The control group yielded similar ANOVA results, with a significant main effect of MD (F(2,22) = 34.058, *p* < 0.001, partial η2 = 0.756; all post-hoc comparisons were significant at *p* < 0.007). Additionally, the control group exhibited a main effect of phase (F(2,22) = 9.359, *p* = 0.003, partial η2 = 0.460) without interaction (F(2,22) = 2.026, *p* = 0.151, partial η2 = 0.156). Post-hoc comparisons revealed that the early phase had a significantly larger area than the middle and late phases (early vs. middle: *p* = 0.004, D = 0.805; early vs. late: *p* = 0.018, D = 0.945). Thus, both groups appear to slightly improve their speed-accuracy trade-off over repeated tests, with no evidence suggesting that spaceflight adversely affects this trade-off.

**Figure 2—figure supplement 4.**
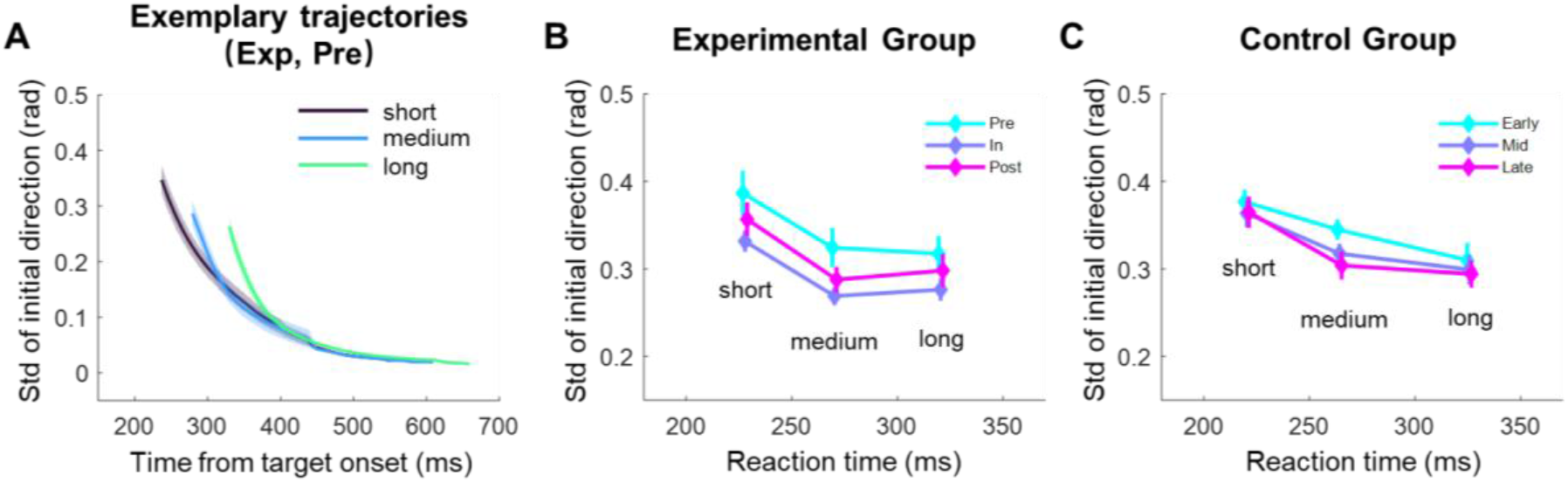
Speed-accuracy trade-off in action planning, illustrated by the relationship between reaction time (RT) and the variance in initial movement direction. **A)** Similar to the speed-accuracy trade-off analysis in Figure S5, all trials were grouped into three equal-sized bins based on RT, regardless of phase, movement direction, or beep condition. The standard deviation of movement direction is shown over time for these bins, using data from the experimental group in the pre-flight phase as an example. The standard deviation decreases over time within a reaching movement, with shorter RTs resulting in greater directional variance, particularly in the early phase of movement. **B-C)** The standard deviation of initial movement direction as a function of RT, plotted separately for the three phases. A negative trend between standard deviation and RT indicates a speed-accuracy trade-off in action planning. However, phase did not affect the relationship between RT and initial variance. For the experimental group, a 3 (phase) × 3 (RT) two-way repeated measures ANOVA on the standard deviation showed a significant main effect of RT (F(2,22) = 7.542, *p* = 0.009, partial η2 = 0.407), while neither the main effect of phase nor the interaction was significant (phase: F(2,22) = 3.160, *p* = 0.083; interaction: F(4,44) = 0.503, *p* = 0.694). Pairwise comparisons indicated that the initial variance of short-RT trials was significantly higher than that of medium-RT trials (*p* = 0.004, D = 0.375), while the difference between medium- and long-RT trials was only marginally significant (*p* = 0.076). The control group showed similar results (RT: F(2,22) = 4.268, *p* = 0.040, partial η2 = 0.280; phase: F(2,22) = 1.657, *p* = 0.215; interaction: F(2,22) = 0.007, *p* = 0.998). Thus, while clear indications of a speed-accuracy trade-off in action planning were observed, there was no evidence that microgravity negatively affected this relationship.

**Figure 2—figure supplement 5.**
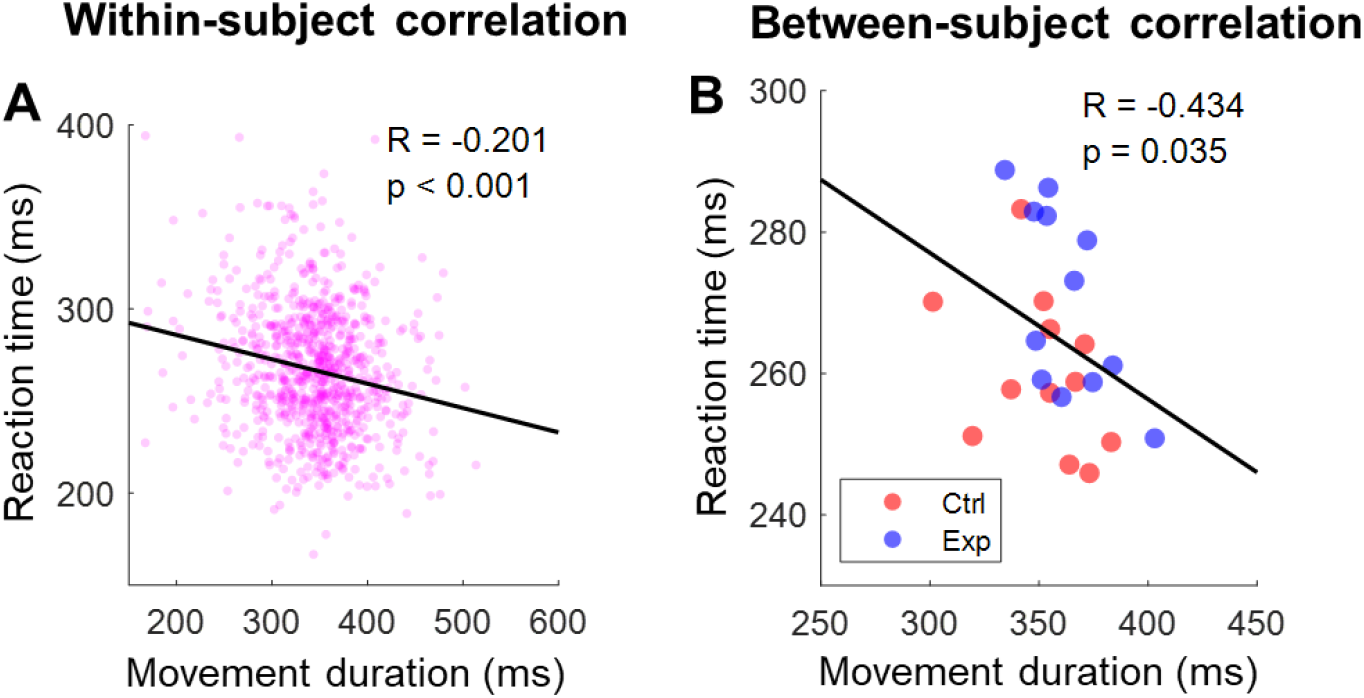
Correlation between reaction time (RT) and movement duration (MD). **A)** Within-subject correlation for a representative participant, whose R value is close to the median of all participants (*R*_*med*_ = −0.230). All trials were pooled, with each pink dot representing the MD and RT of a single reaching movement. The black line indicates the linear fit. Across all 24 participants, 22 showed significant negative correlations between MD and RT, with R values ranging from −0.149 to −0.408 (all p < 0.001). **B)** Between-subject correlation of MD and RT. For each participant, MD and RT were calculated as the mean and median of all trials, respectively. Red and blue dots represent participants in the control and experimental groups, respectively. Across all 24 participants, MD was significantly correlated with RT (r = −0.434, p = 0.035), indicating that, on an individual level, faster movements were associated with slower RTs. Furthermore, no changes in the negative slope between MD and RT were observed across test sessions or phases, suggesting that microgravity does not impact taikonauts’ ability to flexibly adjust their movement timing to meet task demands.

**Figure 2—figure supplement 6.**
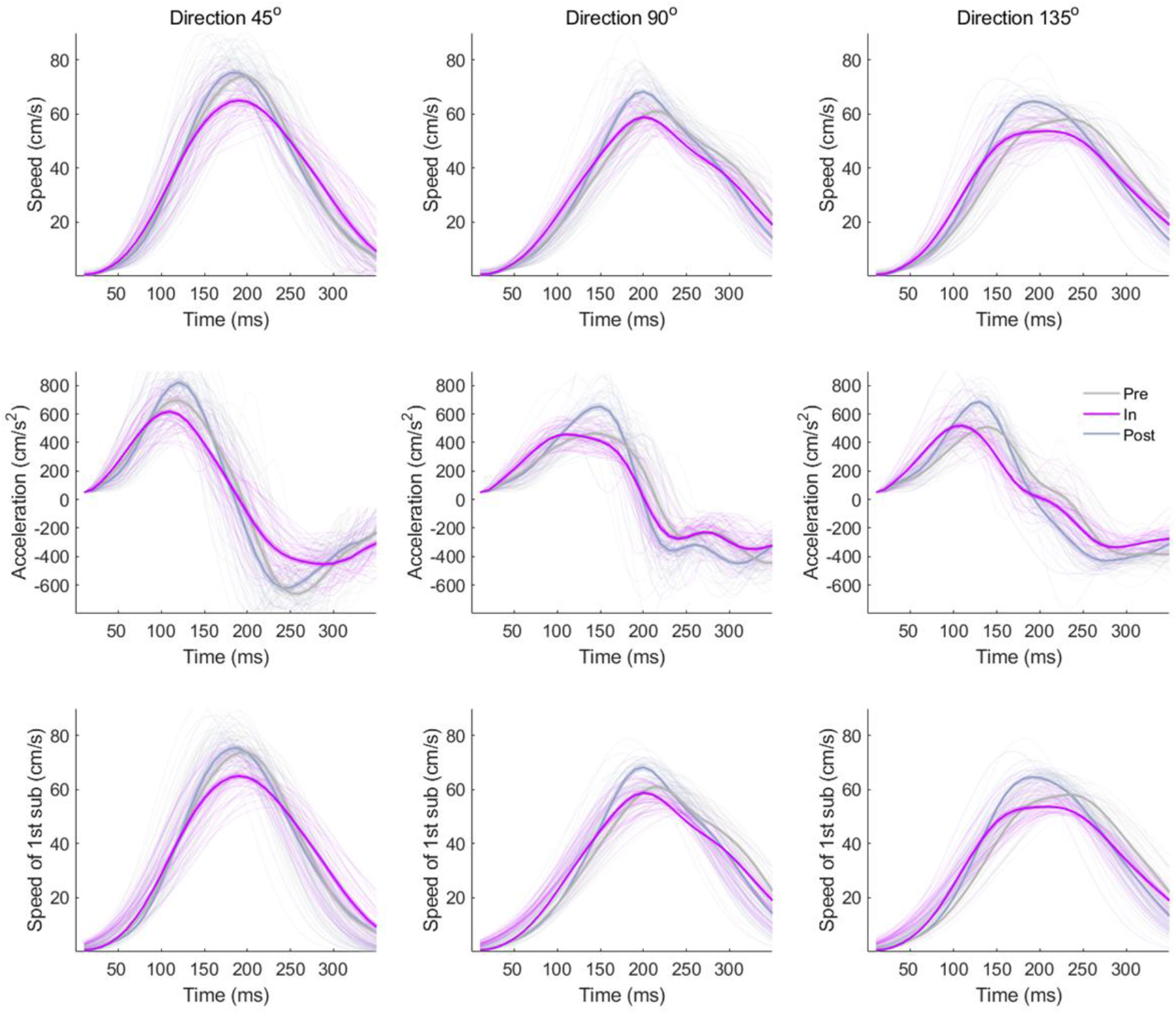
Hand speed profiles (upper panels), hand acceleration profiles (middle panels) and speed profiles of the primary submovements towards different directions from an example participant.

**Figure 3—figure supplement 1.**
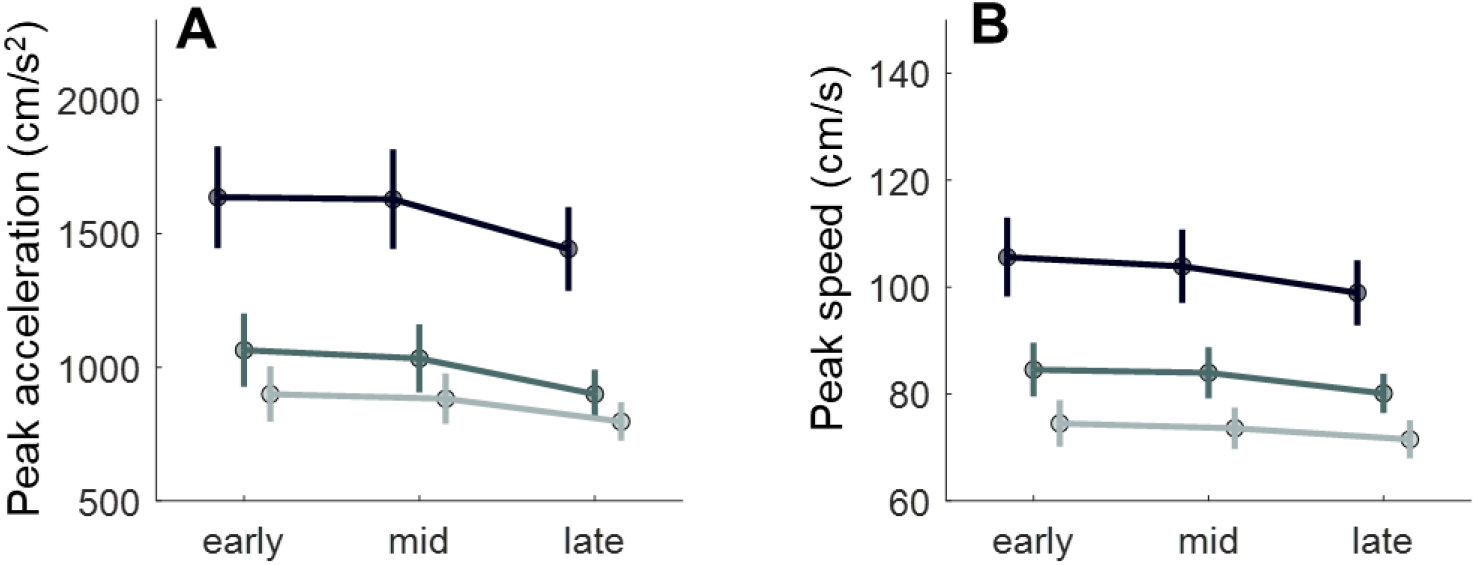
Magnitude changes of peak acceleration and speed of the control group. The control group exhibited no phase effect for peak acceleration and peak speed. Each measure was analyzed using a 3 (direction) × 3 (phase) two-way repeated-measures ANOVA. **A)** Peak acceleration: Significant main effect of direction (F(2,22) = 55.460, *p* < 0.001, partial η2 = 0.834), with all pairwise comparisons significant at p < 0.001. A marginal phase effect (F(2,22) = 3.462, *p* = 0.081) and interaction (F(4,44) = 2.715, *p* = 0.067) were noted. **B)** Peak speed: Significant main effect of direction (F(2,22) = 66.071, *p* < 0.001, partial η2 = 0.857), with all pairwise comparisons significant at p < 0.001. No phase effect (F(2,22) = 2.360, *p* = 0.148) or interaction (F(4,44) = 1.975, *p* = 0.145) was observed.

**Figure 4—figure supplement 1.**
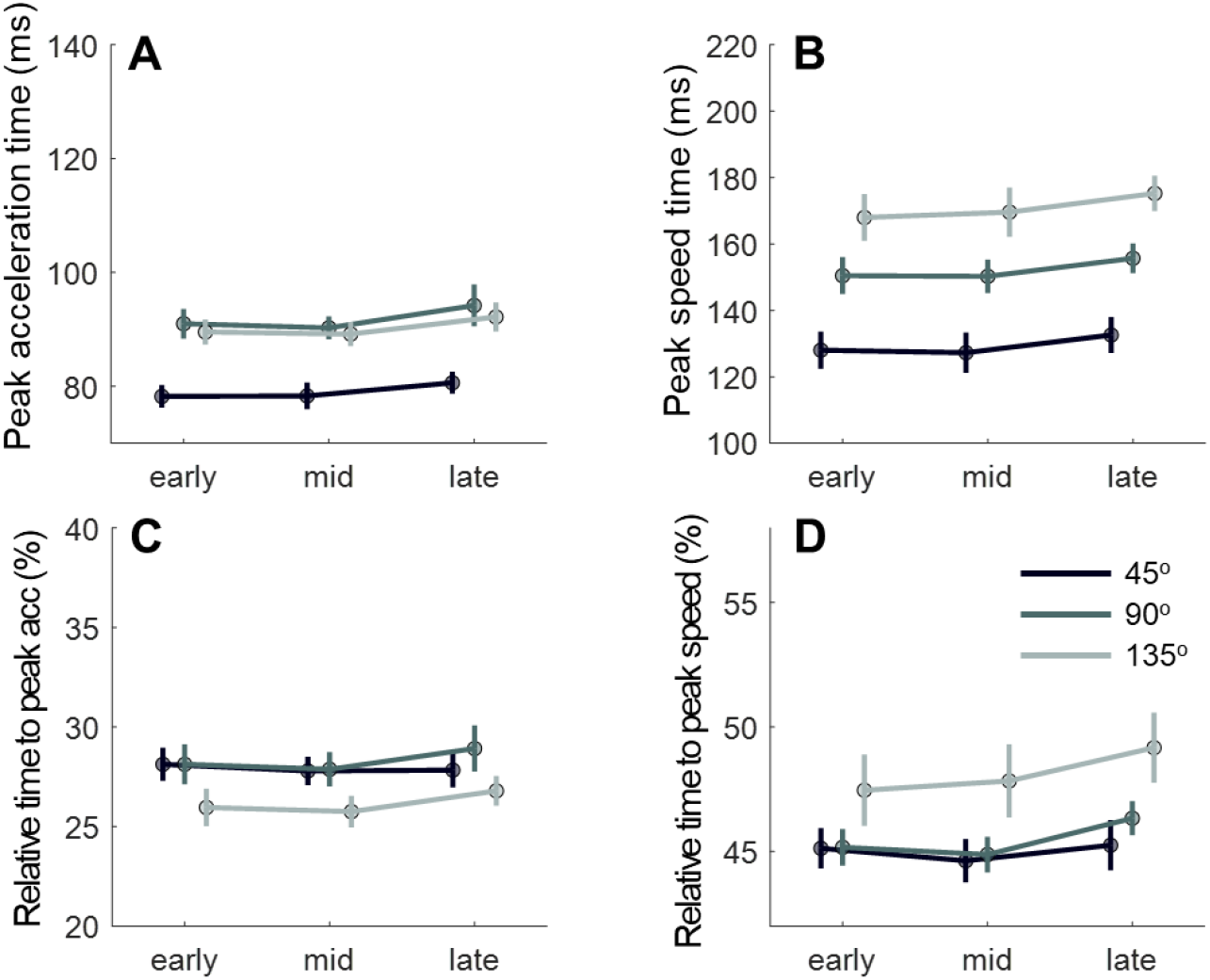
Temporal analysis of peak acceleration and peak speed of the control group. The control group exhibited no phase effect all the performance measures. Each measure was analyzed using a 3 (direction) × 3 (phase) two-way repeated-measures ANOVA. **A)** Peak acceleration time: Significant main effect of direction (F(2,22) = 26.602, *p* < 0.001, partial η2 = 0.707). No phase effect (F(2,22) = 1.440, *p* = 0.260) or interaction (F(4,44) = 0.132, *p* = 0.900) was observed. Pairwise comparisons showed significantly shorter times for 45° compared to 90° (*p* < 0.001, D = 1.574) and 135° (*p* < 0.001, D = 1.386). **B)** Peak speed time: Significant main effect of direction (F(2,22) = 96.575, *p* < 0.001, partial η2 = 0.898), with all pairwise comparisons significant at *p* < 0.001. Marginal phase effect (F(2,22) = 2.702, *p* = 0.098), and no interaction effect (F(4,44) = 0.420, *p* = 0.710). **C)** Relative peak acceleration time: Significant main effect of direction (F(2,22) = 8.506, *p* = 0.003, partial η2 = 0.436), with no phase effect (F(2,22) = 1.143, *p* = 0.317) or interaction (F(4,44) = 1.002, p = 0.381). Pairwise comparisons indicated that relative peak acceleration time appeared significantly earlier in the 135° direction than in 90° (*p* = 0.002, D = 0.913) and 45° (*p* = 0.021, D = 0.747). **D)** Relative peak speed time: Significant main effect of direction (F(2,22) = 6.779, *p* = 0.007, partial η2 = 0.381), with a marginal phase effect (F(2,22) = 2.636, *p* = 0.095) and no interaction (F(4,44) = 1.788, *p* = 0.175). Pairwise comparisons showed that relative peak speed time was significantly earlier in the 135° direction than in 90° (*p* = 0.037, D = 0.686) and 45° (*p* = 0.029, D = 0.803).

**Figure 4—figure supplement 2.**
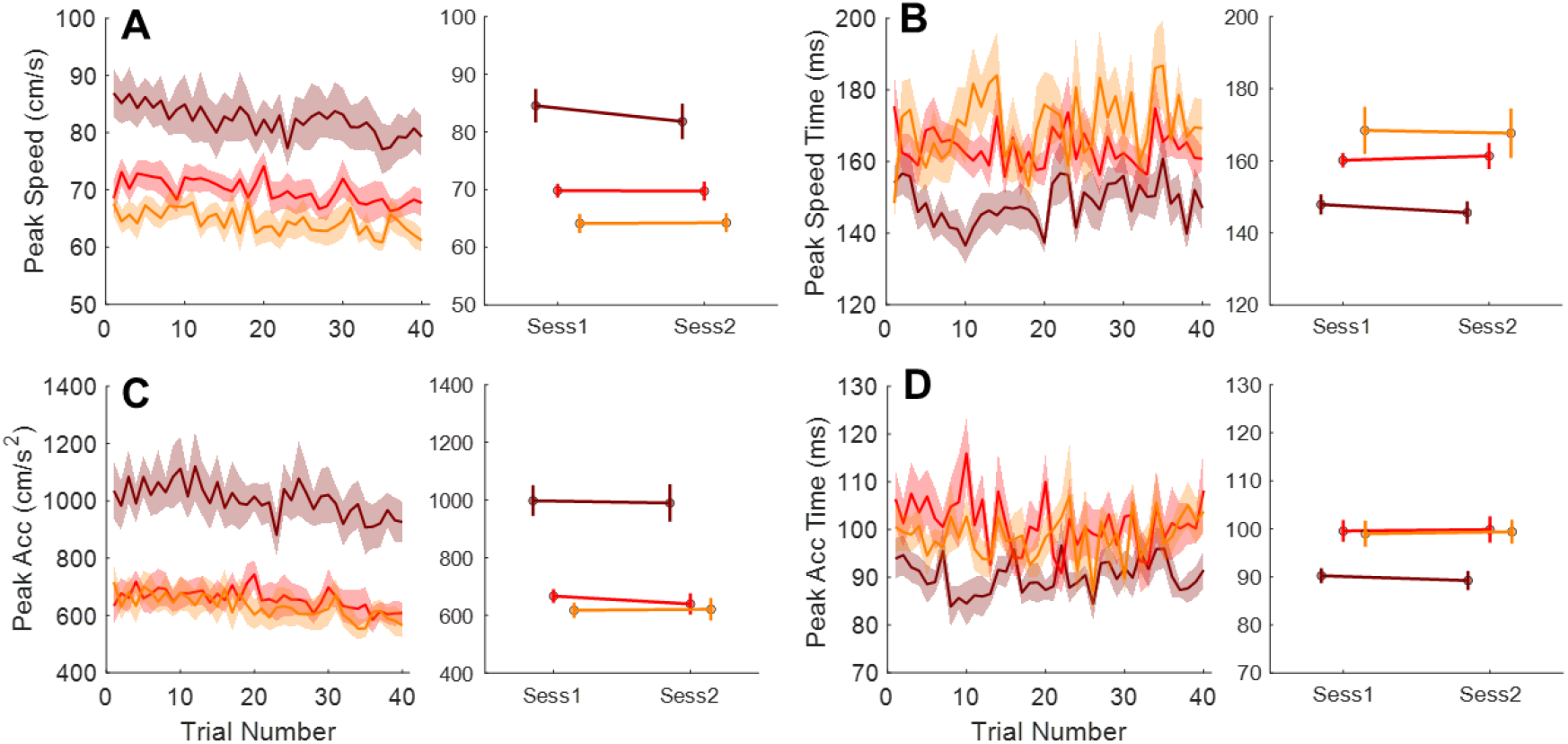
Changes in key performance measures over trials within and between in-flight sessions. To show a lack of within-session and between-session adaptation during spaceflight, we analyzed changes in peak speed **(A)**, peak speed timing of the primary submovement **(B)**, peak acceleration **(C)**, and peak acceleration timing **(D)** across the first and second sessions, as well as trials within the second in-flight session. The second session was specifically analyzed because the sensorimotor system undergoes recalibration during early flight and thus the first session is less likely to show good within-session adaptation. The third session had fewer participants (only six taikonauts). Participant S2, who completed only one in-flight session, was excluded from this analysis. For the within-session analysis, we compared the mean values of the first and last five trials for each measure across movement directions. No significant changes were observed in peak speed timing and peak acceleration timing (all *p*-values > 0.06; panels B and D). Peak speed (panel A) showed a slight and marginally significant decrease in the 45° and 90° directions (45°: from 85.9 ± 3.4 cm/s to 78.5 ± 3.4 cm/s, *p* = 0.013; 90°: from 71.6 ± 2.1 cm/s to 67.3 ± 1.8 cm/s, p = 0.052), but no significant change in the 135° direction (*p* = 0.200). Similarly, peak acceleration (panel C) decreased slightly across all directions with marginal significance (45°: from 1033.9 ± 73.4 cm/s2 to 919.9 ± 66.8 cm/s2, *p* = 0.056; 90°: from 671.8 ± 43.4 cm/s2 to 603.7 ± 33.8 cm/s2, *p* = 0.097; 135°: from 663.2 ± 46.3 cm/s2 to 586.2 ± 36.2 cm/s2, *p* = 0.049). For comparisons between the first and second sessions, we applied a two-way repeated measures ANOVA to these dependent variables and found no significant differences between sessions (all *p*-values > 0.144). These findings suggest that the mass underestimation effects persist over repetitive reaching attempts (120 trials) and practice sessions. Instead of showing between-trial or between-session adaptation, these effects tend to increase slightly across trials.

**Figure 5—figure supplement 1.**
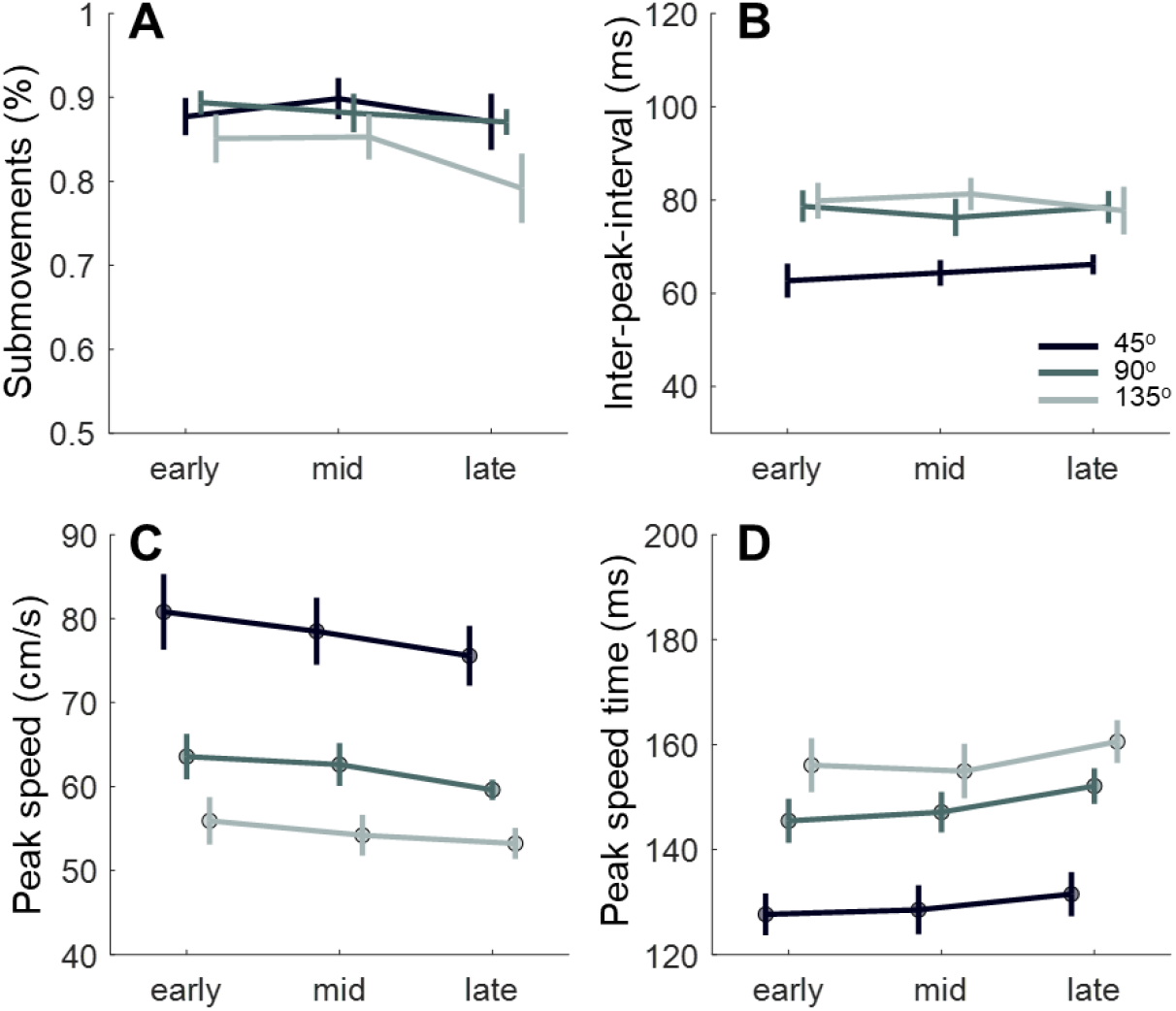
Results of submovement analysis of the control group. All measures of submovements did not show significant phase effect. **A)** Percentage of trials with two submovements. Only a marginal main effect of direction was detected (F(2,22) = 3.297, *p* = 0.057, partial η2 = 0.231) without a phase effect (F(2,22) = 2.175, *p* = 0.159) or interaction (F(4,44) = 1.199, *p* = 0.323). **B)** Inter-peak interval (IPI) showed a significant main effect of direction (F(2,22) = 29.803, *p* < 0.001, partial η2 = 0.730), with no phase effect (F(2,22) = 0.085, *p* = 0.819) or interaction (F(4,44) = 1.559, *p* = 0.202). Pairwise comparisons indicated a significantly shorter IPI in the 45° direction compared to the other two directions (45° vs. 90°: *p* < 0.001, D = 1.180; 45° vs. 135°: *p* = 0.002, D = 1.340). **C)** Peak speed of the primary submovement exhibited a significant main effect of direction (F(2,22) = 55.070, *p* < 0.001, partial η2 = 0.834), without a phase effect (F(2,22) = 3.257, *p* = 0.081) or interaction (F(4,44) = 0.481, *p* = 0.705). Pairwise comparisons showed significant differences between all directions (45° vs. 90°: *p* < 0.001, D = 1.622; 45° vs. 135°: *p* < 0.001, D = 2.429; 90° vs. 135°: *p* < 0.001, D = 0.807). **D)** Peak speed time of the primary submovement revealed a significant main effect of direction (F(2,22) = 43.578, *p* < 0.001, partial η2 = 0.799), with no phase effect (F(2,22) = 2.630, *p* = 0.103) or interaction (F(4,44) = 0.370, *p* = 0.730). Pairwise comparisons showed a significantly smaller peak speed time in the 45° direction compared to the other two directions (45° vs. 90°: *p* < 0.001, D = 1.515; 45° vs. 135°: *p* < 0.001, D = 2.140).

**Table S1.** The schedule of experiment sessions.

| Taikonauts | Pre-flight |  | In-flight |  |  | Post-flight |  |
| --- | --- | --- | --- | --- | --- | --- | --- |
| S1 | -43 | -24 | 39 | 74 | -- | 14 | 66 |
| S2 | -43 | -24 | 39 | -- | -- | 14 | -- |
| S3 | -43 | -24 | 39 | 74 | -- | 14 | 66 |
| S4 | -157 | -60 | 32 | 96 | 131 | 18 | 73 |
| S5 | -165 | -60 | 32 | 96 | 131 | 18 | 73 |
| S6 | -165 | -60 | 32 | 96 | 131 | 18 | 73 |
| S7 | -31 | -12 | 20 | 109 | -- | 15 | 64 |
| S8 | -31 | -12 | 20 | 109 | -- | 15 | 64 |
| S9 | -31 | -12 | 20 | 109 | -- | 15 | 64 |
| S10 | -91 | -48 | 28 | 76 | 125 | 16 | 93 |
| S11 | -91 | -50 | 28 | 77 | 125 | 16 | 93 |
| S12 | -91 | -52 | 28 | 76 | 125 | 16 | 93 |

Numbers in the table represent days before launch (pre-flight), days after launch (in-flight), and days after landing (post-flight).

## Supplementary Note 1

Cost scaling alone cannot explain earlier timing with lower peaks To test whether the kinematic changes observed in microgravity—reduced peak velocity and acceleration together with earlier peak times—can be explained solely by changes in optimal control strategy (i.e., cost-function rescaling) without altering the estimated mass.

We used the same LQG framework as in the main text with cost function:

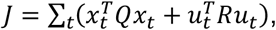

planning a primary movement with fixed nominal distance and duration. The simplified arm model used to derive predictions for direction effects is identical to that in the main text; here we hold the estimated mass constant and vary only the control costs.

Following Crevecoeur et al. (2010), we implemented a scaling parameter α by:

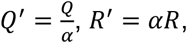

which penalizes control effort more strongly as α increases (more “conservative” strategy), and less strongly as α decreases.

We simulated three movement directions (45°, 90°, 135°) with alpha = 3.0 (increased effort penalty) and alpha = 0.3 (decreased effort penalty). With alpha = 3.0, peak velocity and acceleration decreased but peak times were delayed (Figure 1—figure supplement 2); with alpha = 0.3, peak amplitudes increased and peak times occurred earlier (Figure 1—figure supplement 3). Across directions, no alpha produced the simultaneous combination of lower peak values and earlier peak times that we observed in microgravity. This pattern also mirrors the trade-off in Crevecoeur et al. (2010, Fig. 7B), where lowering peak velocity via cost-function scaling is consistently accompanied by later peaks, not earlier ones. We therefore conclude that changes in optimal-control costs alone (via Q/R rescaling) cannot account for the microgravity pattern of reduced peak velocity/acceleration together with earlier peak times.

## Supplementary Note 2

Coriolis and centripetal torques have minimal impact

To evaluate the potential contribution of Coriolis and centripetal torques, we examined how adding the Coriolis and centripetal torque term to the optimal control simulations affects the outcomes. Using the iLQR (iterative LQR) method of Li & Todorov (2004), we generated OFC-based hand-movement trajectories (Figure R4-R5) and computed the contributions of each torque component separately (Figure R6). Key methods and results are summarized below.

Following Li & Todorov (2004), the forward dynamics of joint angles are given by:

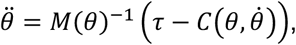

where:

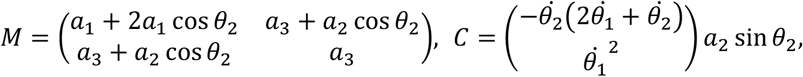

with parameter definitions:

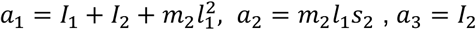

The system is expressed in state-space form:

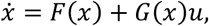

where:

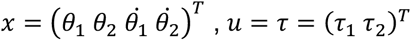

The cost function is defined as:

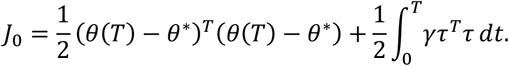

All mechanical parameters and muscle time constants were kept consistent with those used in the main text. Because the dynamics are nonlinear, we solved the OFC problem using the iLQR/iLQG trajectory optimization scheme (Li & Todorov, 2004), which fits a sequence of local LQR subproblems (backward Riccati pass and forward rollout) to obtain a locally optimal time-varying feedback controller for the nonlinear system.

We extracted the hand endpoint position as:

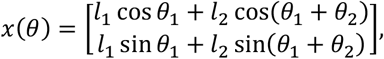

Hand speed and acceleration were computed (Figure 1—figure supplement 4, left panels, solid lines). Using the same utility model to determine movement times, we also simulated hand position, speed, and acceleration under microgravity (dashed lines). The right panels replicate results from the simplified model. Overall, the model with Coriolis and centripetal torques produces similar trajectories to the original, simplified model.

Figure 1—figure supplement 4 compares simulation results from the full model (with Coriolis and centripetal torques, left panels) and the simplified model (right panels). The top row shows hand position profiles, with speed and acceleration shown below. The three colors represent different movement directions (dark red: 45°, red: 90°, yellow: 135°). Dashed lines show simulations under microgravity, assuming mass underestimation. In the full model, a two-link arm was used with both inertial and Coriolis and centripetal torque terms.

These results support our original simplification: excluding Coriolis and centripetal torques does not affect the model’s core predictions about mass underestimation and its effects on motor behavior.

To further evaluate Coriolis and centripetal torque contributions, we followed Hollerbach & Flash (1982, *Biol. Cybern*.) by decomposing net torque τ into inertial and CC components:

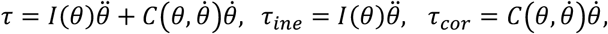

As shown in Figure 1—figure supplement 5, the Coriolis and centripetal term (dashed red) contributes only modestly to the net torque, while the inertial term (dotted red) dominates. This differs from Hollerbach & Flash, likely because their movements involved larger joint excursions and higher peak speeds (≈30–90° vs. <30° here) and larger peak torques (~15 N·m vs. ~4 N·m here). Given the small 12 cm reaches in our paradigm, Coriolis and centripetal torques remain comparatively minor across all three directions, and they had only a negligible effect on overall kinematics, as shown in Fig. R4. We therefore conclude that inertial torques—modulated by mass underestimation—are the primary drivers of movement slowing in microgravity. Misestimation of Coriolis and centripetal torques cannot explain our empirical findings, since Coriolis and centripetal torques play only a minor role in this context.

